# embarcadero: Species distribution modelling with Bayesian additive regression trees in R

**DOI:** 10.1101/774604

**Authors:** Colin J. Carlson

## Abstract

1. `embarcadero` is an `R` package of convenience tools for species distribution modelling with Bayesian additive regression trees (BART), a powerful machine learning approach that has been rarely applied to ecological problems.
2. Like other classification and regression tree methods, BART estimates the probability of a binary outcome based on a set of decision trees. Unlike other methods, BART iteratively generates sets of trees based on a set of priors about tree structure and nodes, and builds a posterior distribution of estimated classification probabilities. So far, BARTs have yet to be applied to species distribution modelling.
3. `embarcadero` is a workflow wrapper for BART species distribution models, and includes functionality for easy spatial prediction, an automated variable selection procedure, several types of partial dependence visualization, and other tools for ecological application. The `embarcadero` package is available open source on Github and intended for eventual CRAN release.
4. To show how `embarcadero` can be used by ecologists, I illustrate a BART workflow for a virtual species distribution model. The supplement includes a more advanced vignette showing how BART can be used for mapping disease transmission risk, using the example of Crimean-Congo haemorrhagic fever in Africa.

## 1 Introduction

In the last two decades, over two dozen statistical and machine learning methods have been proposed for species distribution modelling (SDM) (Norberg *et al.*, 2019). Over time, a handful of methods have risen to predominance due to ease of implementation, computational speed, and strong predictive performance in rigorous cross-validation. Some methods are especially popular for specific applications, mostly because of disciplinary tradition. For example, maximum entropy (MaxEnt) models are widely popular for studies of global ecological responses to climate change (VanDerWal *et al.*, 2013; Warren *et al.*, 2013). In disease ecology, boosted regression trees (BRTs) have become the dominant tool for mapping vectors, reservoirs, and transmission risk of infectious zoonoses and vector-borne diseases (Carlson *et al.*, 2019; Pigott *et al.*, 2014; Messina et al., 2016), largely due to an influential 2013 paper on dengue virus (Bhatt et al., 2013). SDMs are used for several—sometimes conflicting—purposes in ecology, and popular methods are sometimes used despite known shortcomings (Guillera-Arroita et al., 2015; Smith & Santos, 2019). In particular, most popular methods have a limited framework for handling uncertainty, and conspicuously few popular methods are Bayesian (and vice versa).

In this paper, I discuss a new Bayesian approach to classification and regression trees (CART), one of the most popular families of machine learning methods used in ecology. Models in this family estimate the probability of a given output variable (in this case, a binary classification of habitat suitability or species presence) based on decision “trees” that split predictor variables with nested, binary rule-sets. The precise rules for generating these trees vary across implementations. For example, in *random forest* models, an ensemble of trees is generated, where each tree is independently generated based on a boostrap of the original dataset; trees grow to the maximum possible depth (the longest chain of splitting rules), with no pruning (trees are never *post hoc* reduced). In the *boosted regression trees* approach (BRT), shallower trees with a constrained depth (“weak learners”) are constructed iteratively that explain the residuals left by previous trees; this adds bias, but allows the model to focus on unusual cases at the potential expense of overfitting (Elith *et al.*, 2008; Vezhnevets & Barinova, 2007). CART methods have many strengths for species distribution modelling; they consistently perform well in model comparisons (Elith *et al.*, 2006; Mainali *et al.*, 2015; Redding *et al.*, 2017; Wisz *et al.*, 2008), and the tree-based approach is often more intuitive than the complex fitting procedures “under the hood” of MaxEnt or Maxlike methods (Elith *et al.*, 2011; Merow *et al.*, 2013; Merow & Silander, 2014).

*Bayesian additive regression trees* (BART) are an exciting and new alternative to other popular classification tree methods. As in other approaches, BART generates a set of decision trees that explain different components of variance in the outcome variable. Unlike random forests or boosted regression trees, the formulation of BART is Bayesian, with the posterior probability of a model shaped by priors *P*(trees) on how trees should look (i.e., the parameters used to generate those trees):

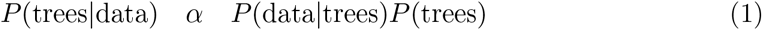

Like boosted regression trees, BART introduces variance by fitting a set of many shallow “weak learner” trees, but unlike BRT, this is explicitly controlled by three prior distributions: the probability a tree stops at a node of a given depth, the probability of a given variable being drawn for a splitting rule, and the probability of splitting that variable at a particular value. The latter two are usually treated as uniformly distributed (splits are randomly constructed by variable, and within each variables’ range), while the first is usually specified as a negative power law, constraining tree depth and penalizing overfitting. Using these priors, a specified number of trees *m* are generated with no splits, and then updated randomly in an MCMC process that allows them to be expanded, rearranged, or pruned. Each model instance is a *sum-of-trees* model, unlike random forests, which average predictions across trees; to create the sum-of-trees model, each tree is adjusted to the residuals of the sum-of-remaining-trees. This process superficially resembles how boosting works within boosted regression trees, but because trees are tuned to the ensemble, they rarely overfit to particular cases within the residuals. (Chipman *et al.*, 2010) After dropping a burn-in period, the full set of sum-of-trees models from different points in the Markov chain is treated as a posterior distribution, and used to generate the posterior distribution of predictions. (For a more in-depth explanation, including a visualization of tree structure in the MCMC process, see Tan & Roy (2019).)

In computer science, BARTs are used for everything from medical diagnostics to selfdriving car algorithms (Sparapani *et al.*, 2018; Tan *et al.*, 2018); however, they have yet to find any widespread application in ecology. A study from 2011 used BART as a tool to examine habitat selection data on birds (Yen *et al.*, 2011); a 2017 study used BART to evaluate performance data of other species distribution modelling methods (Farley, 2017). But so far, they have not been used for the purpose of predicting species distributions. This reflects a broader deficit of Bayesian models in the SDM literature: several elegant Bayesian SDM methods have been previously proposed (Golding & Purse, 2016; Redding *et al.*, 2017), but none are particularly widely adopted, possibly because advanced Bayesian models may seem discouraging or unintuitive.

BART brings the conceptual familiarity and strengths of classification tree methods, but adds a relatively simple Bayesian component that inherently and intuitively handles model uncertainty. This might make it a promising alternative not just to existing Bayesian approaches but also popular classification tree methods, in particular boosted regression trees. BRT has several easy to use out-of-the-box implementations, is powerful for ecological inference, and consistently performs well in rigorous tests of SDM performance. However, BRT also has downsides: it can be prone to overfitting, and fitting procedures are largely handed down as anecdotal best practices, with many studies choosing hyperparameters based on software defaults; very few studies select parameters from formal cross-validation as early work recommended (Elith *et al.*, 2008). Furthermore, uncertainty is usually measured by generating an unweighted ensemble of BRT submodels over subsetted training data, generating a confidence interval from data permutations (like random forests) rather than formal assumptions about model uncertainty. In contrast, the formal Bayesian structure of BART captures uncertainty within a single model, which is more coherent and intuitive than how uncertainty is usually generated in BRT ensembles. BART also shares many of the strengths of BRT, like easy out-of-the-box implementation and easy visualization of “black box” model components, and outperforms other CART methods in model comparisons. (Chipman *et al.*, 2010)

This paper introduces an `R` package, `embarcadero`, as a convenience tool for running SDMs with BARTs. Throughout, I use a simulated “virtual species” (see Appendix 1) to illustrate the workflow and the major features of the package, including model selection, visualization, and diagnostics. Because boosted regression trees are the most popular method of species distribution modelling in medical geography, the supplement includes a second, more detailed vignette using BART to map Crimean-Congo haemorrhagic fever (CCHF) in Africa, based on the distribution of the tick *Hyalomma truncatum*, a presumed vector. This is a more challenging and computationally-intensive implementation, and takes several hours to run on most machines, but highlights some of the strength of the approach for applied scientific questions.

## 2 SDMs with BARTs

### 2.1 Implementing BART with binary classification

At least four `R` packages currently exist that can implement BARTs: `BayesTree` (Chipman & McCulloch, 2016), `bartMachine` (Kapelner & Bleich, 2013), `BART` (McCulloch *et al.*, 2018), and `dbarts` (Chipman *et al.*, 2014). Their functionality differs in important ways, and not all of them are currently capable of important features like partial dependence plots that are important for SDMs. This package is an SDM-oriented workflow wrapper for `dbarts`, which includes most of the basic functionality needed for species distribution modelling, including a simple implementation of BART with binary outcomes. A list of the functions made available in `embarcadero`, versus their counterparts and additional useful functions in `dbarts`, is given in Table 1.

**Table 1:** Functions available in `embarcadero` and additional functions in `dbarts` of importance.

| Core modelling functionality |  |
| --- | --- |
| `bart` (in `dbarts`) | Runs a binary BART classification model. |
| `bart.step` | Full implementation of a BART model with built-in variable set reduction (a wrapper for `dbarts:::bart`, `variable.step`, `varimp`, `varimp.diag`, and `summary`). |
| `predict` | Predict species distributions with a BART model and a `RasterStack` of environmental layers (a wrapper for `dbarts:::predict.bart`). |
| `summary` | Returns a summary of call, performance, and diagnostic plots for a BART model object. |
| Variable diagnostics |  |
| `variable.step` | Stepwise variable set reduction algorithm. |
| `varimp` | Returns variable importance, with optional plots. |
| `varimp.diag` | Diagnostic of variable importance at different $m$ values. |
| Visualization |  |
| `partial` | Partial dependence plots for single variables (a `ggplot2`-based wrapper for `dbarts:::pdbart`). |
| `pd2bart` (in `dbarts`) | Two-predictor, three-dimensional partial dependence plots (no wrapper implemented yet). |
| `plot.mcmc` | Visualize each posterior draw's prediction and the running average of those predictions. Can be used with the `animation` package to create GIFs of how the posterior draw learns to fit the data (especially interesting for the burn-in of models with small number of trees). |
| `spartial` | Spatial projection (maps) of partial dependence plots onto raw environmental covariates. |
| Convenience tools |  |
| `bigstack` | Fast aggregation of an environmental layer `RasterStack` for quick prediction, using the `velox` package. |

In the original notation of Chipman *et al.* (2010), BART consists of tree structures *T* and terminal nodes (leaves) *M*, as an ensemble (*T*_1_, *M*_1_),…, (*T_n_*, *M_n_*). Each tree generates a predictive function *g*(·), with a sum of trees function *f* (·) given as

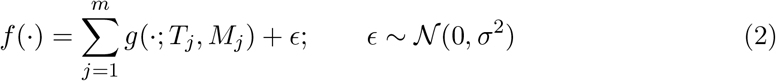

A set of posterior draws of *f*\*, generated by the MCMC process described above, create the posterior distribution for *p*(*f*|*y*) ≡ *p*(trees\data). Given the assumption of normality, BART handles binary classification problems (like species distribution modelling) using a logit link, where Φ is the standard normal c.d.f. and:

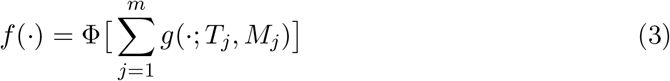

Binary classification is run by `dbarts::bart` automatically when supplied with a binary outcome. However, the returned predictions are untransformed back into probabilities, a problem solved in `embarcadero` with a `predict` wrapper. (This also allows prediction on raster datasets, a key piece of SDM workflow.)

### 2.2 An example of a BART SDM

To see how BART works, we can generate a virtual species on a hypothetical landscape which responds to climate variables X1 through X4, but is uninfluenced by variables X5 to X8 (see Appendix 1). Like most other SDM methods in R, the BART model itself is run on a data frame of presence-absence or presence-pseudoabsence points, and associated environmental covariates. For example, with a `RasterStack` called `climate` and an occurrence dataset called `occ`.df, the basic workflow is

~~~
library(embarcadero)
xnames <- c(‘x1, ‘x2’, ‘x3’, ‘x4’,
               ‘x5’, ‘x6’, ‘x7’, ‘x8’)
## Run the BART model
sdm <- bart(y.train=occ.df[,‘bserved’],
                   x.train=occ.df[,xnames],
                   keeptrees = TRUE)
## Predict the species distribution
map <- predict(sdm, climate
## Visualize model performance
summary(bart)
~~~

This last line returns a brief model diagnostic including the optimal cutoff for thresholding classifications and some measures of performance, like the area under the receiveroperator curve (AUC):

~~~
Call: bart occ.df[, xnames] occ.df[, “Observed”] TRUE
Predictor list:
   x1 x2 x3 x4 x5 x6 x7 x8
Area under the receiver-operator curve
   AUC =0.91
Recommended threshold (maximizes true skill statistic)
   Cutoff = 0.42
   TSS = 0.71
   Resulting type I error rate: 0.078
   Resulting type II error rate: 0.21
~~~

Additionally, summary returns a diagnostic figure (**Figure 1**), summarizing the performance of the classifier on the training data.

**Figure 1:**
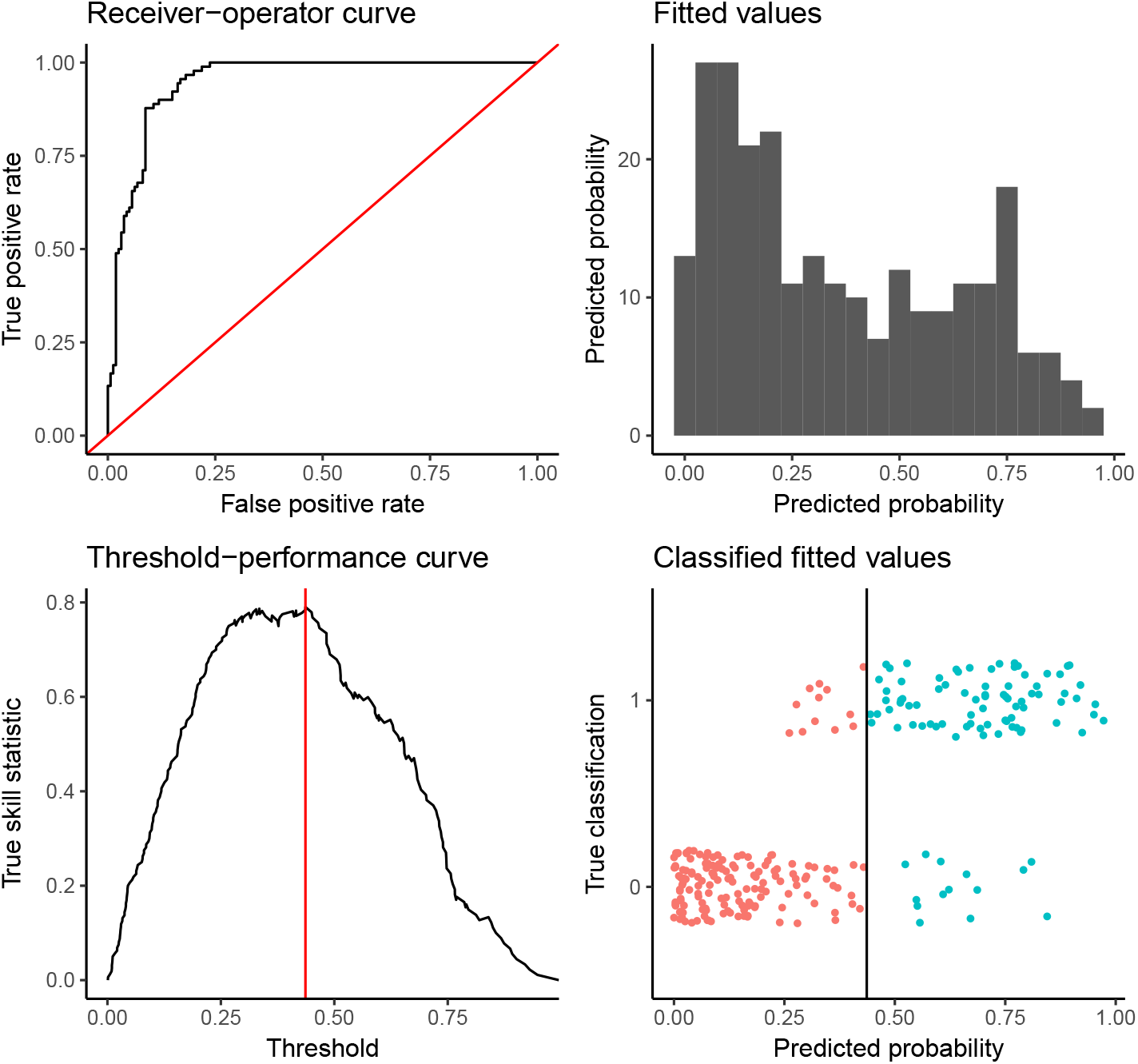
The model diagnostic returned by `summary()`.

The primary appeal of BART, compared to other CART methods, is a formal way of measuring model uncertainty within any individual implementation. Pulling uncertainty out of BART predictions is easy with embarcadero; for example, to pull a 95% credible interval, a user can specify:

~~~
map <- predict(sdm, climate, quantiles=c(0.025, 0.975))
~~~

Mapping the difference between these two rasters gives the credible interval width, which provides a native measure of spatial uncertainty, analogous to how the coefficient of variation can be used to measure spatial uncertainty across an ensemble of BRT runs (Carlson *et al.*, 2019). When running tasks especially with several quantiles, or large rasters, prediction runtime grows quickly and memory can become limiting; predict() has a “splitby” option that breaks the task into pieces, which minimizes memory conflicts, adds a progress bar, and allows estimation of total runtime based on the first chunk:

~~~
map <- predict(sdm, climate, quantiles=c(0.025, 0.975), splitby=10)
~~~

## 3 Variable selection

Variable importance (calculated by `varimp()`) is usually measured in BART models by counting the number of times a given variable is used by a tree split across the full posterior draw of trees. (This is similar to variable importance in BRTs, which is calculated from the number of tree splits and the corresponding improvement they cause in the model.) In models with higher numbers of trees, the difference in variable importance becomes less pronounced, and less informative variables receive a higher number of splitting rules. Conversely, variable selection can be performed by running models with a small number of trees (*m* = 10 or 20), and observing which variables stop being included in trees. (Chipman *et al.*, 2010) This diagnostic is generated in `embarcadero` by `varimp.diag()` (see an example in **Figure 2**).

**Figure 2:**
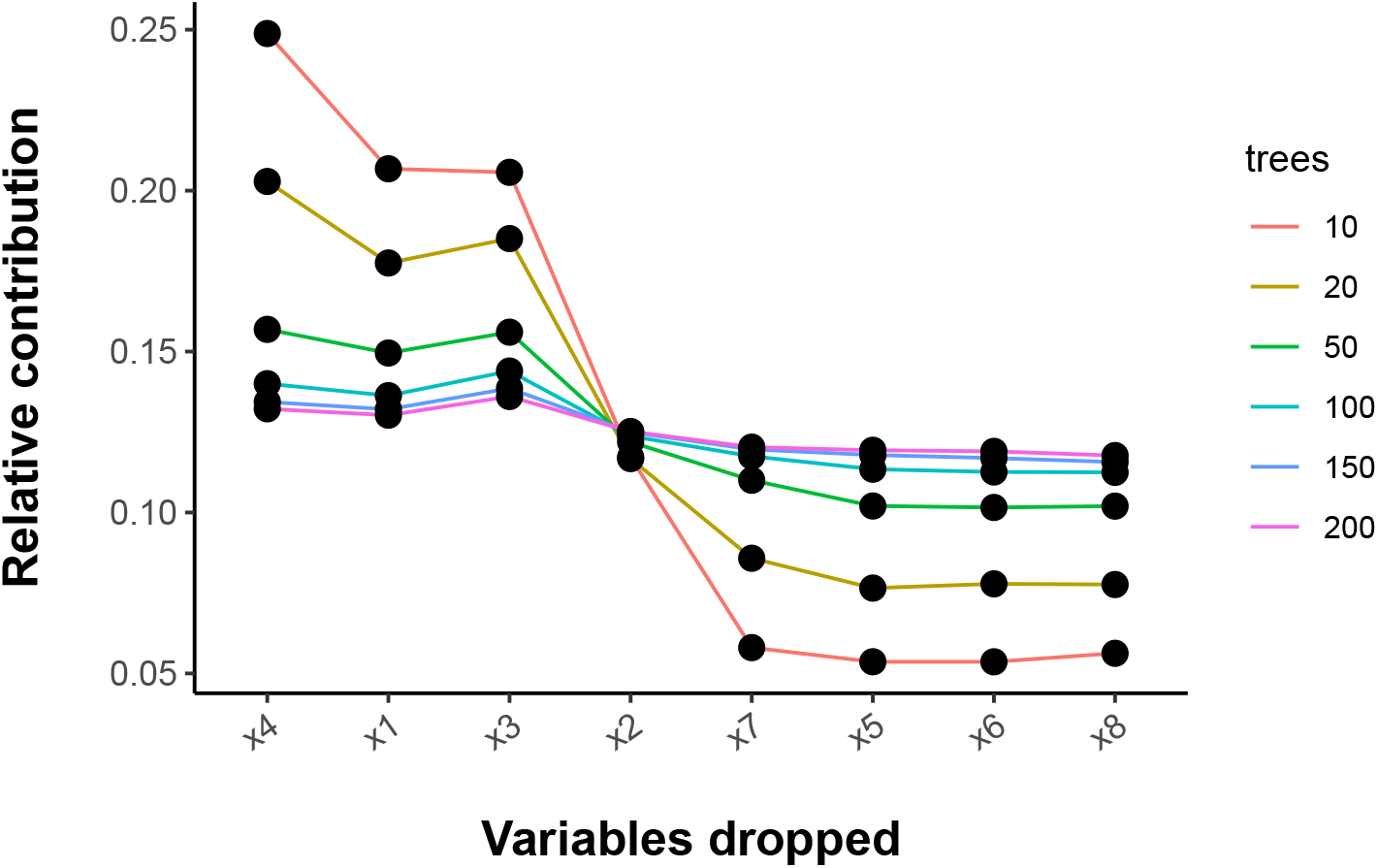
The model diagnostic returned by `varimp.diag()`.

Analysis of this diagnostic plot is still subjective and informal. As a way to standardize variable set reduction rules across workflows, `embarcadero` includes an automatic variable selection procedure in `variable.step():`

i. Fit a full model with all predictors and a small tree ensemble (default *m* = 10), a fixed number of times (default *n* = 50)
ii. Eliminate the least informative variable across all 50 runs;
iii. Re-run the models again minus the least informative variable (*n* = 50 times again), recording the root mean square error (on the training data);
iv. Repeat steps 2 and 3 until there are only three covariates left;
v. Finally, select the model with the lowest average root mean square error (RMSE).

Anecdotally, this procedure almost always recommends dropping every variable with decreasing importance in models with fewer trees, and conserves every variable with increasing importance. In our virtual species case, for example, the diagnostic shows that X1 through X4 have much higher performance than X5 through X8 (**Figure 2**), and the automated procedure recommends dropping X5 through X8:

~~~
varimp.diag(occ.df[,xnames],
       occ.df[,”Observed”],
       iter=50)
step.model <- variable.step(x.data=occ.df[,xnames],
                    y.data=occ.df[,”Observed”])
step.model
[1] “x1” “x2” “x3” “x4”
~~~

This largely matches original work which found that BART is highly effective at identifying informative subsets of predictors (see section 5.2.1 of Chipman *et al.*, 2010).

I recommend careful analysis of all diagnostic information, but include a full automated variable selection pipeline in `bart.step`, which (a) produces the initial multi-*m* diagnostic plot, (b) runs automated variable selection, (c) returns a model trained with the optimal variable set, (d) plots variable importance in the final model, and (e) returns the summary of the final model. Despite automation, this procedure is not a fail-safe against the inclusion of uninformative predictors, or false inference on them; this is true of almost all methods, and predictors should always be chosen based on at least some expert opinion about biological plausibility (Fourcade *et al.*, 2018). Similarly, validation of partial depencence curves against biological knowledge should be treated as an additional level of model validation, potentially more informative than measuring predictive accuracy (Warren *et al.*, 2019).

## 4 Visualizing model results

`embarcadero` includes several methods for generating partial dependence plots. The function `partial` is written as a wrapper for `dbarts::pdbart`, and can be used to generate partial dependence plots with a customizable, `ggplot2`-based aesthetic, including multiple ways of visualizing uncertainty. (As with overall predictions, credible intervals on partial plots are true Bayesian credible intervals.) Posteriors can be visualized with traceplots of individual draws, or bars for a credible interval of a specified width (by default 95%):

~~~
partial(sdm, x.vars=c(“x4”),
              smooth=5,
              equal=TRUE,
              trace=FALSE)
## VERSUS, for comparison,
gbm1 <- dismo::gbm.step(data=occ.df,
                       gbm.x = 2:5, gbm.y = 1,
                       family = “bernoulli”,
                       tree.complexity = 5,
                       learning.rate = 0.01,
                       bag.fraction =0.5)
dismo::gbm.plot(gbm1, variable.no=4, rug=TRUE,
                          plot.layout=c(1,1))
~~~

This visualizes uncertainty much clearer than, for example, `dismo::gbm.plot` can in a single instance (**Figure 3**). Two-dimensional partial dependence plots (interactions among two predictor variables) can also be generated using `dbarts::pd2bart`.

**Figure 3:**
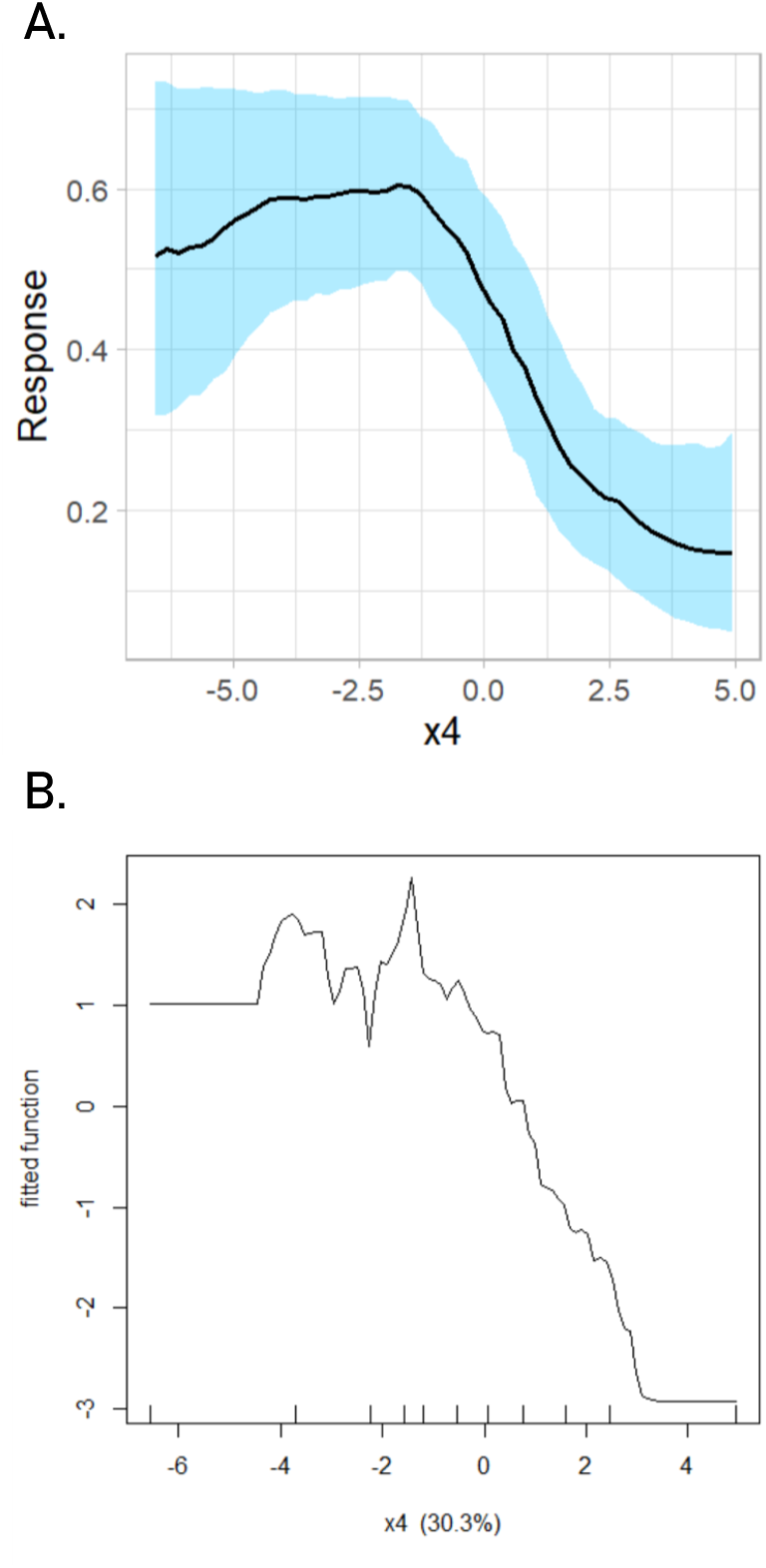
Partial dependence curves generated by single-instance BART implementations (A) show uncertainty with more transparency and clarity than those generated from single-instance BRT implementations (B).

Finally, `embarcadero` a new visualization called *spatial partial dependence plots*, which reclassify predictor rasters based on their partial dependence plots, and show the relative suitability of different regions for an individual covariate. The `spartial` function can be used to generate these maps, and answer questions like “What desert regions are too arid, even in their wettest month, for spadefoot toads?” or “Where are the soils with the best pH for redwood growth?” These visualization options are illustrated in greater depth in the advanced vignette.

## 5 An advanced vignette

To demonstrate applications to disease transmission mapping, the supplement includes an advanced tutorial on embarcadero focused on updating an African risk map for Crimean-Congo haemorrhagic fever virus (CCHF). CCHF is a tick-borne Bunyavirus that causes extremely severe, and often fatal, illness in humans. Very little is known about CCHF, compared to other cosmopolitan tick-borne illnesses like Lyme disease or tularemia. The definitive reservoir of CCHF is unknown but likely ungulates (Babayan *et al.*, 2018); outbreaks frequently affect sheep and other domestic ruminants. The vectors of CCHF are better known, and are presumed to almost always be *Hyalomma* ticks, which are widespread throughout Africa and Eurasia; other tick vectors have been suspected, but evidence for their competence is limited. (Papa *et al.*, 2017) In Africa, *Hyalomma trun-catum* in particular is common throughout rangeland and is a strong candidate for a primary vector. (Logan *et al.*, 1989; Wilson *et al.*, 1991) A global map of Crimean-Congo haemorrhagic fever has been previously been produced with boosted regression trees; a significant amount of the Black Sea region was suitable, while areas outside had highly localized predictions of suitability, presumably because of data sparsity in Africa especially. (Messina *et al.*, 2015b) However, some major areas of presence appeared under-predicted, such as the western Congo Basin.

The advanced vignette shows how BART can be used to map CCHF in Africa, using the same occurrence dataset as previous mapping efforts have (Messina *et al.*, 2015a). Just as studies of dengue risk have included suitability for the *Aedes aegypti* mosquito as a covariate, the new model includes a suitability layer for *Hyalomma truncatum*, created from the canonical dataset on African tick distributions. (Cumming, 1998). The updated map predicts that the distribution of CCHF may be more geographically expansive than previous studies have indicated (**Figure 4**). Areas of the highest risk are still heavily concentrated in Sahel rangeland and east African highlands, but also far more extensive in southern Africa and along the Atlantic coast than previously believed. A detailed tutorial is provided showing this workflow in the Supplementary Materials of this paper, and all data are available online (github.com/cjcarlson/pier39).

**Figure 4:**
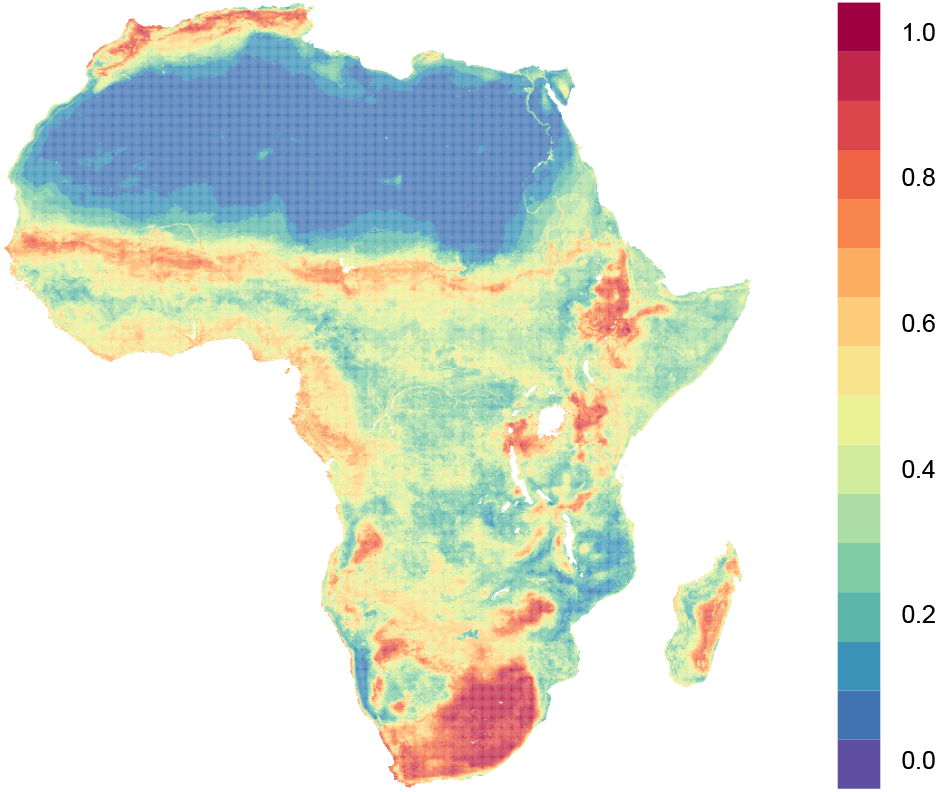
A map of Crimean-Congo haemorrhagic fever transmission risk, constructed using ecological niche modelling with BART (see Supplementary Materials).

## 6 Discussion

Because BART is a comparatively new method, many of the basic use case questions remain mostly unaddressed: Do pseudoabsences perform notably worse than absences? Is there a minimum sample size? Does collinearity inflate or distort variable importance? Users may wish to explore some of these points using virtual species before working with BART on their data, or to compare BART results to other methods as a sense check.

Furthermore, as with any other Bayesian method, out of the box implementation can make it easy to neglect or underconsider prior selection. More advanced users may be interested in going more in depth within the BART literature to set better priors. For example, using a uniform prior on covariate importance can be unhelpful—especially in high-dimensionality data with only a few valid predictors, where the model tends to converge on the variable importance prior. (Tan *et al.*, 2018; Rocková & van der Pas, 2017) Instead, setting a Dirichlet distribution for the prior can significantly improve model performance and variable selection. (Linero, 2018)

Finally, it is worth mentioning that BART is a growing topic of interest in machine learning, and new extensions may expand applications within SDM work and more broady in spatial ecology. For example, the random intercept BART (riBART) model is a framework for handling cases of structure within outcome data; this framework might be useful for cases where sampling bias has categorical structure (e.g., different levels of sampling across country or state borders). (Tan *et al.*, 2018) Similarly, causal inference using the BART framework has become especially popular (Hahn *et al.*, 2017), which may be an interesting direction for modelling given recent work proposing causal inference as a new priority for mapping infectious diseases. (Kraemer *et al.*, 2019) Expanding work along these lines will help establish better best practices for using BARTs in SDM applications.

## Supporting information

Supporting Information: Advanced Vignette

## Acknowledgements

Thanks to Vincent Dorie for creating the fabulous and wonderfully useful `dbarts` package, to Shweta Bansal and Jason Blackburn for research support, to Ethan Beaman for telling me about BARTs in the first place, to Zack Susswein for helpful comments on design of Bayesian models, to Tad Dallas for helpful comments on the R package and manuscript, and to David Lawrence Miller and a second anonymous reviewer for detailed and helpful feedback. This work was funded by a Georgetown Environment Initiative (GEI) postdoctoral fellowship.

## Appendix 1. Generating a virtual species for modelling

For this example, we create a virtual landscape of eight Gaussian “climate variables” on a 150 by 150 cell grid (with `NLMR`), create a virtual species inhabiting that landscape but only depending on four of eight total “climate variables” (with `virtualspecies`), and then extract a presence-absence dataset for modelling (with `embarcadero`).

~~~
library(NLMR, quietly = T)
library(virtualspecies, quietly = T)
set.seed(12345)
~~~

~~~
## Random landscape
onelandscape <- function(x) {NLMR::nlm_gaussianfield(nrow = 150,
                                             ncol = 150,
                                             rescale = FALSE)}
climate <- stack(lapply(c(1:8), onelandscape))
names(climate) <- c(“x1”,”x2”,”x3”,”x4”,”x5”,”x6”,”x7”,”x8”)
## Generate the species’ climatic niche from X1 through X4
random.sp <- generateRandomSp(climate[[1:4]],
                     approach=“pca”,
                     relations=“gaussian”,
                     species.prevalence=0.5,
                     realistic.sp = TRUE,
                     PA.method=“threshold”)
## Generate some presences, and some absences
sp.points <- sample0ccurrences(random.sp,
                    n=250,
                    type = “presence-absence”)
## Extract the associated climate values
occ <- SpatialPoints(sp.points$sample.points[,c(“x”,”y”)])
occ.df <- cbind(sp.points$sample.points,
                raster::extract(climate, occ))
## Finally, let’s drop the long-lats and the “Real” presence-absence
## values and just leave behind an “Observed” and the climate data occ.df <- occ.df[,-c(1:3)]
~~~

If we were to run head(occ.df) it should return a data frame that looks like this:

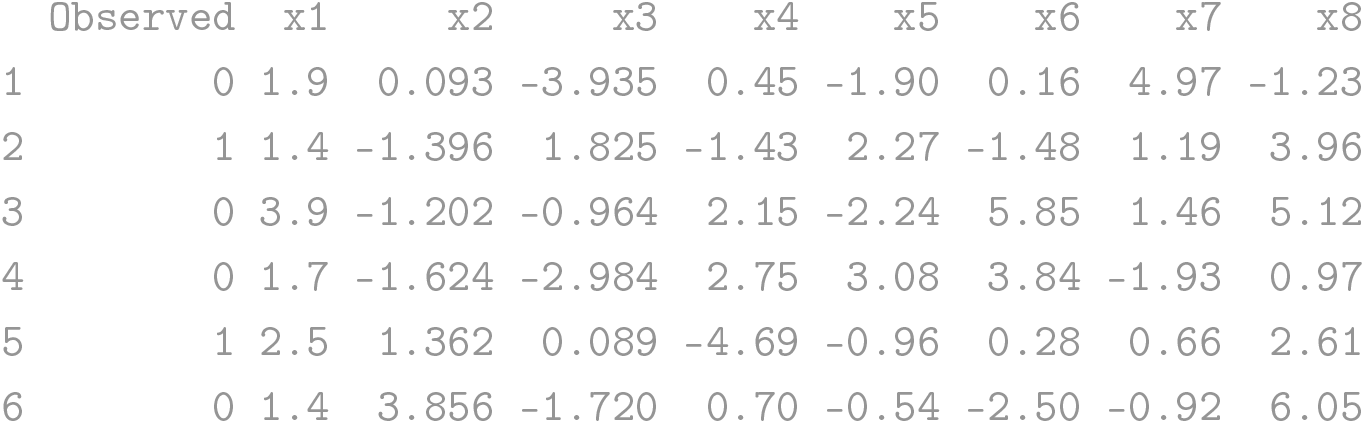

