## Supporting Information: Advanced Vignette for "embarcadero: Species distribution modelling with Bayesian additive regression trees in R"

### Mapping Crimean-Congo Haemorrhagic Fever in Africa with BART

#### Getting Started

So you're interested in using `embarcadero` to do species distribution modeling with Bayesian additive regression trees! That's great. BARTs are a powerful way to do machine learning and, while not a new method per se, they are very new for SDMs.

In this advanced tutorial, I'm going to assume that you've seen the main paper and the `embarcadero` package vignette, which goes over basic functions and some of the internal structure of BART models. This tutorial, and the associated repo (`cjcarlson/pier39`), are intended for advanced users interested in seeing what a professional workflow might look like for an applied use.

```
library(embarcadero)
set.seed(12345)
```

Our goal here is to do a few things:

1. Build a species distribution model for *Hyalomma truncatum*, a tick we think might be a vector of Crimean-Congo haemorrhagic fever (CCHF) in Africa.
2. To build a transmission risk map, again using species distribution modeling/ecological niche modeling, for CCHF in Africa.
3. To see if we learn anything from that model about CCHF we didn't know before.
4. To see some of the advanced visualization and workflow tricks in the `embarcadero` package!

#### Mapping Hyalomma

We're going to make a suitability layer for *Hyalomma truncatum*, a possible CCHF vector. The tick occurrence data, which is a set of presence points, comes from Cumming et al. (1998) *Bulletin of Entomological Research*, one of the most detailed datasets in the world on parasite distributions. We're going to build a species distribution model using those points, some climate data, and a few other convenient layers that might be relevant.

##### Data entry

Let's start by loading in the climate data. We'll use three main sources:

1. WorldClim v1.4 (Hijmans et al. 2005), a harmonized dataset of bioclimatic variables mostly used for species distribution modeling
2. Two layers describing long-term averages of the normalized difference vegetation index (NDVI), which captures the greenness of a landscape (taken from previous use in Carlson et al. 2018). This can be an important variable for measuring invertebrate distributions, for example, or partitioning landscapes into different biomes.
3. A layer produced by the NASA SEDAC center describing the percent cropland by grid cell. If we think ruminants are spreading CCHF, this might matter! (https://sedac.ciesin.columbia.edu/data/set/aglands-croplands-2000)

Based on some expert opinion, I picked a handful of variables I thought might work well (and it's already reduced down to minimize redunandancy):

- BIO1: Mean annual temperature
- BIO2: Mean diurnal range
- BIO5: Max temperature of warmest month
- BIO6: Min temperature of coldest month
- BIO12: Mean annual precipitation
- BIO13: Precipitation of wettest month
- BIO14: Precipitation of driest month
- BIO15: Precipitation seasonality
- Mean NDVI (normalized difference vegetation index)
- NDVI amplitude
- Percent cropland by pixel

Let's read in the covariates.

```
load(file = "~/Github/pier39/covsraw.rda"); covs <- covsraw
class(covs)
#> [1] "RasterStack"
#> attr(,"package")
#> [1] "raster"
covs@crs <- crs('+proj=longlat +ellps=WGS84 +datum=WGS84 +no_defs +towgs84=0,0,0')
```

Next, let's load in our tick occurrence dataset, which is a set of longitude/latitude coordinates.

```
load(file = "~/Github/pier39/ticks.rda")
head(ticks)
#>       X     ScientificName Longitude.X Latitude.Y
#> 1 16471 Hyalomma truncatum          16       -8.2
#> 2 16472 Hyalomma truncatum          14      -12.5
#> 3 16473 Hyalomma truncatum          12       -5.5
#> 4 16474 Hyalomma truncatum          15      -16.5
#> 5 16475 Hyalomma truncatum          14      -13.6
#> 6 16476 Hyalomma truncatum          15      -12.4
nrow(ticks)
#> [1] 1794
```

Presence points are usually a little aggregated in space, so as a normal data cleaning practice, I like to thin my occurrence data to one point per raster grid cell.

```
# Make a spatial point data frame
mod <- SpatialPointsDataFrame(ticks[,3:4],data.frame(ticks[,1]))
names(mod@data) <- 'Presence'
nrow(mod)
#> [1] 1794

# Rasterizing and converting back to points makes one unique point per grid cell
tmp=rasterize(mod, covs[[1]], field="Presence", fun="min")
pts.sp1=rasterToPoints(tmp, fun=function(x){x>0})
nrow(pts.sp1)
#> [1] 1716

# Extract the climate data at each point
pres.cov <- raster::extract(covs, pts.sp1[,1:2])
head(pres.cov)
#>      bio1  bio12  bio13  bio14 bio15 bio2 bio5 bio6  crop ndvi.amp
#> [1,]   29 0.0064 0.0090 0.0042  0.15   23   47  8.0 0.000    0.050
#> [2,]   30 0.0061 0.0092 0.0037  0.17   23   48  8.7 0.000    0.056
```

```
#> [3,]   32 0.0090 0.0117 0.0064  0.16   22   48 11.3 0.000    0.105
#> [4,]   32 0.0085 0.0145 0.0036  0.38   21   47 14.7 0.031    0.032
#> [5,]   31 0.0081 0.0154 0.0028  0.45   21   46 12.4 0.011    0.014
#> [6,]   32 0.0087 0.0148 0.0037  0.38   21   47 14.9 0.044    0.151
#>      ndvi.mean
#> [1,]    -0.015
#> [2,]     0.016
#> [3,]     0.095
#> [4,]     0.127
#> [5,]     0.122
#> [6,]    -0.039
```

Next, let's generate an equal number of pseudoabsences around Africa to the number of presences we have. BARTs are like BRTs in that they are sensitive to assumed prevalence; anecdotally, I strongly suggest using an equal number of presences and absences in your training data. You can experiment with the demo data by changing "nrow(ticks)" to "5000" below if you want to see some weirder model behavior.

```
#Generate the data
absence <- randomPoints(covs,nrow(ticks))
#> Warning in .couldBeLonLat(x, warnings = warnings): CRS is NA. Assuming it
#> is longitude/latitude
abs.cov <- raster::extract(covs, absence)

#Code the response
pres.cov <- data.frame(pres.cov); pres.cov$tick <- 1
abs.cov <- data.frame(abs.cov); abs.cov$tick <- 0

# And one to bind them
all.cov <- rbind(pres.cov, abs.cov)
head(all.cov)
#>   bio1  bio12  bio13  bio14 bio15 bio2 bio5 bio6  crop ndvi.amp ndvi.mean
#> 1   29 0.0064 0.0090 0.0042  0.15   23   47  8.0 0.000    0.050    -0.015
#> 2   30 0.0061 0.0092 0.0037  0.17   23   48  8.7 0.000    0.056     0.016
#> 3   32 0.0090 0.0117 0.0064  0.16   22   48 11.3 0.000    0.105     0.095
#> 4   32 0.0085 0.0145 0.0036  0.38   21   47 14.7 0.031    0.032     0.127
#> 5   31 0.0081 0.0154 0.0028  0.45   21   46 12.4 0.011    0.014     0.122
#> 6   32 0.0087 0.0148 0.0037  0.38   21   47 14.9 0.044    0.151    -0.039
#>   tick
#> 1    1
#> 2    1
#> 3    1
#> 4    1
#> 5    1
#> 6    1

# Let's just clean it up a little bit - remove any missing data
all.cov <- all.cov[complete.cases(all.cov),]
```

Now we have a dataset ready to do some modeling!

#### Running models with dbarts

We could try something really simple on defaults, right out the gate. The `bart` function in `dbarts` can just be run on defaults:

```r
xvars <- names(all.cov)[!(names(all.cov)=='tick')]
xvars
#>  [1] "bio1"      "bio12"     "bio13"     "bio14"     "bio15"
#>  [6] "bio2"      "bio5"      "bio6"      "crop"      "ndvi.amp"
#> [11] "ndvi.mean"
first.model <- bart(all.cov[,xvars], all.cov[,'tick'], keeptrees=TRUE)
#>
#> Running BART with binary y
#>
#> number of trees: 200
#> number of chains: 1, number of threads 1
#> Prior:
#>  k: 2.000000
#>  power and base for tree prior: 2.000000 0.950000
#>  use quantiles for rule cut points: false
#> data:
#>  number of training observations: 3478
#>  number of test observations: 0
#>  number of explanatory variables: 11
#>
#> Cutoff rules c in x<=c vs x>c
#> Number of cutoffs: (var: number of possible c):
#> (1: 100) (2: 100) (3: 100) (4: 100) (5: 100)
#> (6: 100) (7: 100) (8: 100) (9: 100) (10: 100)
#> (11: 100)
#>
#> offsets:
#>  reg : 0.00 0.00 0.00 0.00 0.00
#> Running mcmc loop:
#> iteration: 100 (of 1000)
#> iteration: 200 (of 1000)
#> iteration: 300 (of 1000)
#> iteration: 400 (of 1000)
#> iteration: 500 (of 1000)
#> iteration: 600 (of 1000)
#> iteration: 700 (of 1000)
#> iteration: 800 (of 1000)
#> iteration: 900 (of 1000)
#> iteration: 1000 (of 1000)
#> total seconds in loop: 10.667233
#>
#> Tree sizes, last iteration:
#> [1] 2 2 2 2 3 3 2 3 2 2 2 2 3 3 4 3 3 2
#> 3 3 4 1 4 2 2 3 2 4 3 5 5 2 3 2 3 2 3 1
#> 3 2 1 3 2 2 3 3 3 3 3 4 2 2 2 3 2 2 2 2
#> 2 3 1 2 2 4 2 2 2 2 1 2 3 3 2 3 2 3 2 2
#> 3 2 2 1 2 2 3 2 3 2 4 2 2 2 2 2 3 1 2 3
#> 4 1 3 2 1 2 2 2 3 2 2 2 3 2 4 2 2 2 3 4
#> 3 2 2 2 2 3 2 3 2 4 2 3 2 4 2 2 3 3 2 2
#> 2 2 2 3 2 3 2 1 2 2 3 2 3 2 2 4 3 2 3 4
#> 2 3 2 2 2 2 2 2 2 2 2 2 2 2 3 2 2 2 2 4
#> 3 2 2 3 2 2 3 2 2 2 3 3 3 3 3 2 2 2 3 1
#> 2 3
```

```
#>
#> Variable Usage, last iteration (var:count):
#> (1: 32) (2: 30) (3: 24) (4: 22) (5: 34)
#> (6: 18) (7: 27) (8: 30) (9: 22) (10: 18)
#> (11: 28)
#> DONE BART
```

In reality, we know we want to do variable set reduction, and tune the model a bit from there. In the main vignette you can see the different component parts of that process. Here, we're going to use our most powerful shortcut: `bart.step`. Inside `bart.step`, five things happen:

1. The variable importance diagnostic from `varimp.diag` is generated
2. The stepwise variable set reduction in `variable.step` is run
3. The final model is run with the reduced predictor set
4. The variable importance plot from `varimp` is generated
5. The model summary from `summary` is returned

This is slow but very easy to simply fire off if you're running a large workflow (you can speed it up, in a demo, by reducing iter.step and even more so by reducing iter.plot). Let's do it:

```
sdm <- bart.step(x.data = all.cov[,xvars],
                 y.data = all.cov[,'tick'],
                 full = TRUE,
                 quiet = TRUE)
#>
#>  10 tree models: 100 iterations
#>
#>  20 tree models: 100 iterations
#>
#>  50 tree models: 100 iterations
#>
#>  100 tree models: 100 iterations
#>
#>  150 tree models: 100 iterations
#>
#>  200 tree models: 100 iterations
```

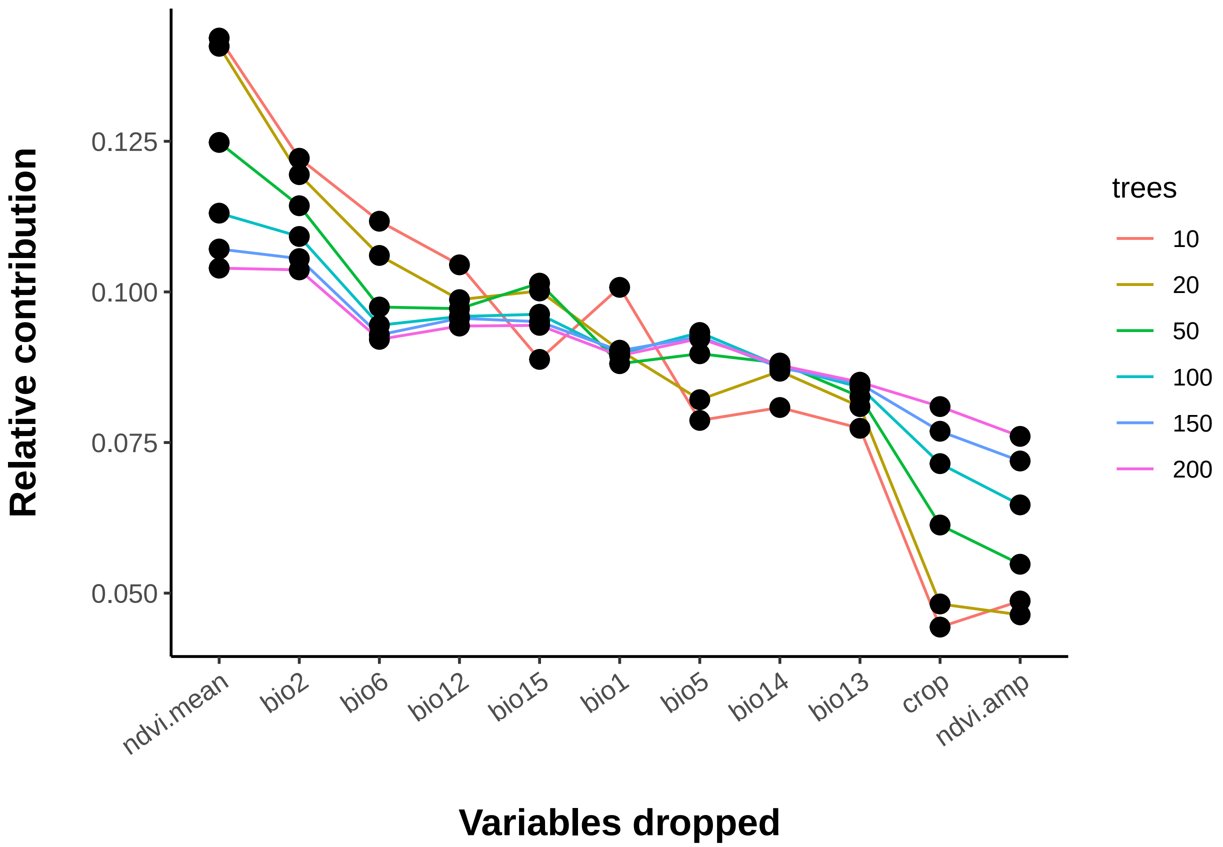


```
#> [1] Number of variables included: 11
#> [1] Dropped:
#> [1]
#> [1] ---------------------------------------
#> [1] Number of variables included: 10
#> [1] Dropped:
#> [1] crop
#> [1] ---------------------------------------
#> [1] Number of variables included: 9
#> [1] Dropped:
#> [1] crop     ndvi.amp
#> [1] ---------------------------------------
#> [1] Number of variables included: 8
#> [1] Dropped:
#> [1] crop     ndvi.amp bio5
#> [1] ---------------------------------------
#> [1] Number of variables included: 7
#> [1] Dropped:
#> [1] crop     ndvi.amp bio5     bio14
#> [1] ---------------------------------------
#> [1] Number of variables included: 6
#> [1] Dropped:
#> [1] crop     ndvi.amp bio5     bio14    bio13
#> [1] ---------------------------------------
#> [1] Number of variables included: 5
#> [1] Dropped:
```

```
#> [1] crop     ndvi.amp bio5     bio14    bio13    bio15
#> [1] ---------------------------------------
#> [1] Number of variables included: 4
#> [1] Dropped:
#> [1] crop     ndvi.amp bio5     bio14    bio13    bio15    bio6
#> [1] ---------------------------------------
#> [1] Number of variables included: 3
#> [1] Dropped:
#> [1] crop     ndvi.amp bio5     bio14    bio13    bio15    bio6     bio2
#> [1] ---------------------------------------
```

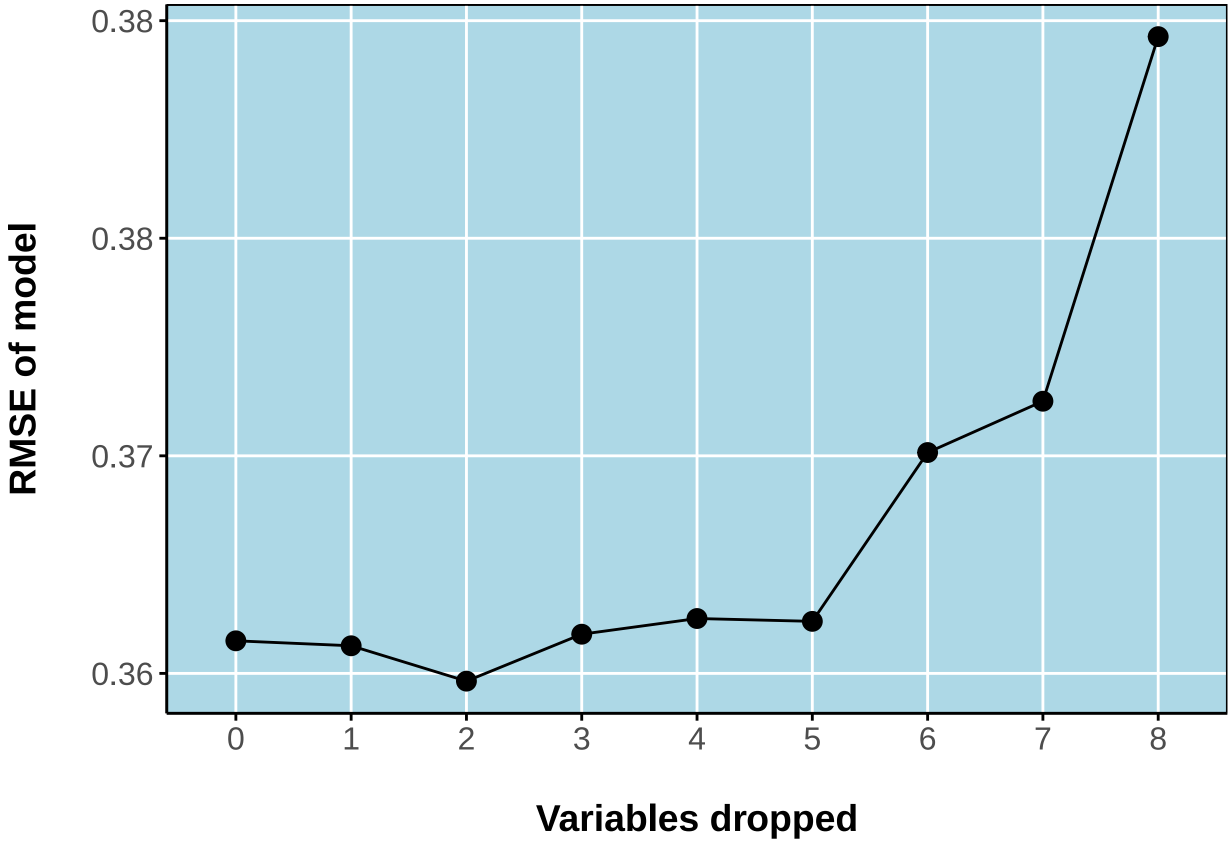


```
#> [1] ---------------------------------------
#> [1] Final recommended variable list
#> [1] bio1      bio12     bio13     bio14     bio15     bio2      bio5
#> [8] bio6      ndvi.mean
#>
#> Running BART with binary y
#>
#> number of trees: 200
#> number of chains: 1, number of threads 1
#> Prior:
#>  k: 2.000000
#>  power and base for tree prior: 2.000000 0.950000
#>  use quantiles for rule cut points: false
#> data:
#>  number of training observations: 3478
#>  number of test observations: 0
```

```
#>  number of explanatory variables: 9
#>
#> Cutoff rules c in x<=c vs x>c
#> Number of cutoffs: (var: number of possible c):
#> (1: 100) (2: 100) (3: 100) (4: 100) (5: 100)
#> (6: 100) (7: 100) (8: 100) (9: 100)
#>
#> offsets:
#>  reg : 0.00 0.00 0.00 0.00 0.00
#> Running mcmc loop:
#> iteration: 100 (of 1000)
#> iteration: 200 (of 1000)
#> iteration: 300 (of 1000)
#> iteration: 400 (of 1000)
#> iteration: 500 (of 1000)
#> iteration: 600 (of 1000)
#> iteration: 700 (of 1000)
#> iteration: 800 (of 1000)
#> iteration: 900 (of 1000)
#> iteration: 1000 (of 1000)
#> total seconds in loop: 8.798700
#>
#> Tree sizes, last iteration:
#> [1] 2 2 2 2 2 3 2 2 1 2 3 1 3 3 3 2 2 2
#> 2 2 4 2 2 4 3 2 2 2 4 2 3 2 2 3 3 2 2 3
#> 2 2 2 3 3 3 3 1 2 4 2 3 3 2 2 2 3 2 2 2
#> 3 2 3 2 2 2 3 2 2 3 2 3 2 2 2 2 2 2 2 3
#> 2 2 3 2 3 2 4 2 3 2 3 3 2 2 2 3 2 3 3 3
#> 3 3 2 6 2 1 2 2 2 2 2 2 2 3 1 3 2 1 2 4
#> 3 3 2 2 2 3 2 2 3 3 2 2 2 2 3 2 2 2 3 2
#> 3 2 4 3 2 2 3 2 2 3 3 3 5 2 2 3 2 2 2 2
#> 2 2 5 3 3 2 4 4 2 2 4 2 3 3 2 2 2 2 2 2
#> 2 2 1 2 2 3 2 2 4 3 2 2 2 3 3 2 2 2 2 2
#> 4 3
#>
#> Variable Usage, last iteration (var:count):
#> (1: 31) (2: 37) (3: 30) (4: 27) (5: 33)
#> (6: 33) (7: 26) (8: 30) (9: 39)
#> DONE BART
```

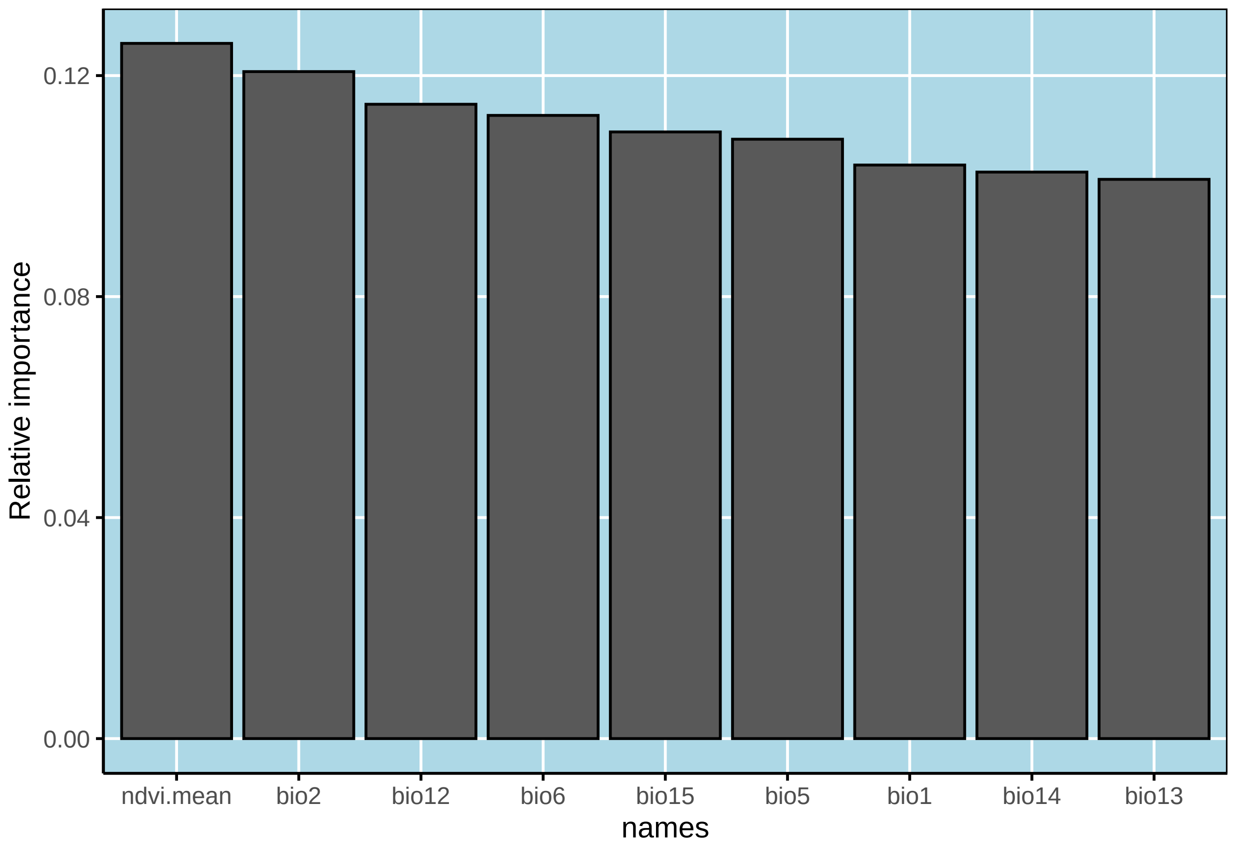


```
#> Call:  bart x.data[, vs] y.data TRUE
#>
#> Predictor list:
#>  bio1 bio12 bio13 bio14 bio15 bio2 bio5 bio6 ndvi.mean
#>
#> Area under the receiver-operator curve
#> AUC = 0.91
#>
#> Recommended threshold (maximizes true skill statistic)
#> Cutoff =  0.53
#> TSS =  0.66
#> Resulting type I error rate:  0.16
#> Resulting type II error rate:  0.19
```

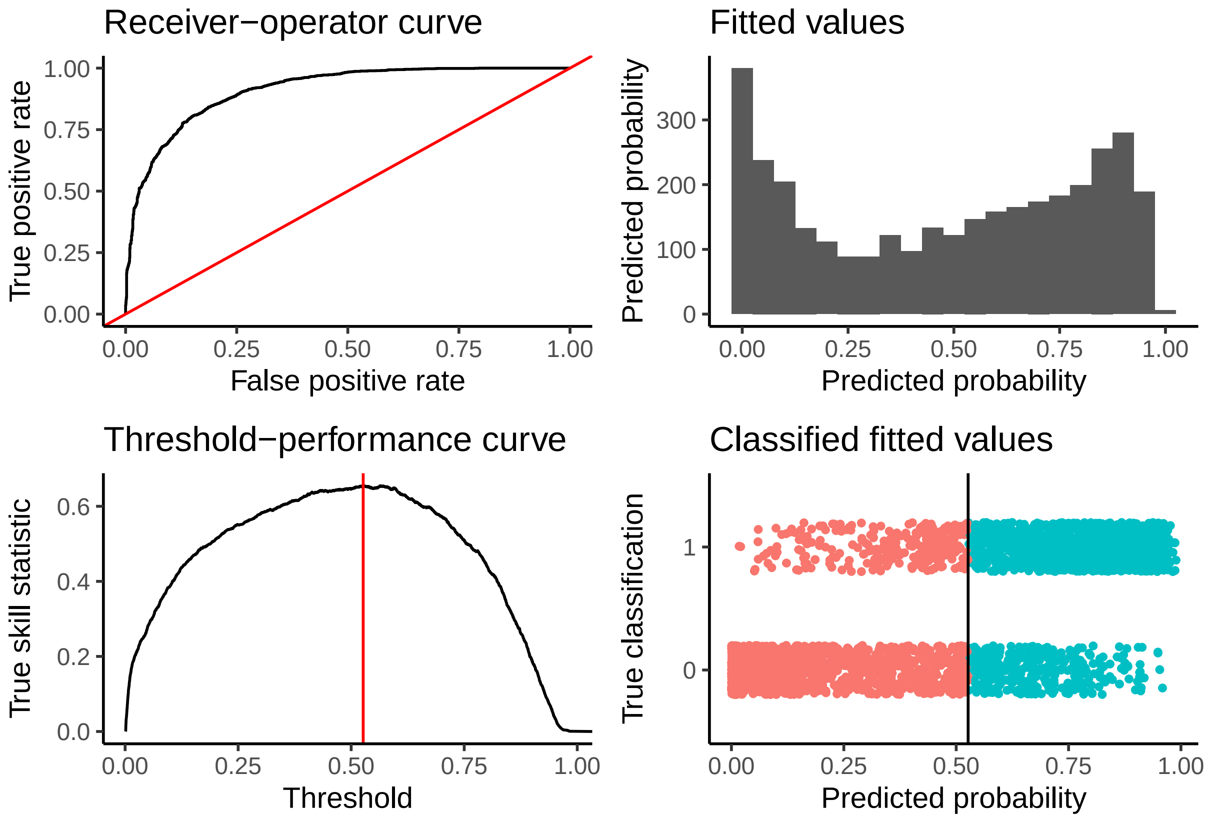


Out of all the variables, "crop" seems to be especially undesirable as a predictor. A few other variables seem like they might not be helping the model either, but we probably need a systematic way to deal with that.

A high AUC value indicates our model performs well. The summary function also returns an optimal threshold that maximizes the true skill statistic (TSS), and the sensitivity/specificity of the model at that cutoff (alpha).

What do the predictions look like? To make a predicted raster, we have to use `embarcadero`'s wrapper for the native `predict` function in `dbarts`.

```
# Do the spatial prediction
hytr.layer <- predict(object = sdm,
                      inputstack = covs,
                      splitby = 20,
                      quiet = TRUE)
#> Estimated time to total prediction (mins):
#> 12

# How's it look?
plot(hytr.layer, box=FALSE, axes=FALSE, main='Hyalomma truncatum')

# Check that against the presence data:
points(SpatialPoints(ticks[,c('Longitude.X','Latitude.Y')]),
       pch=16, cex=0.2)
```

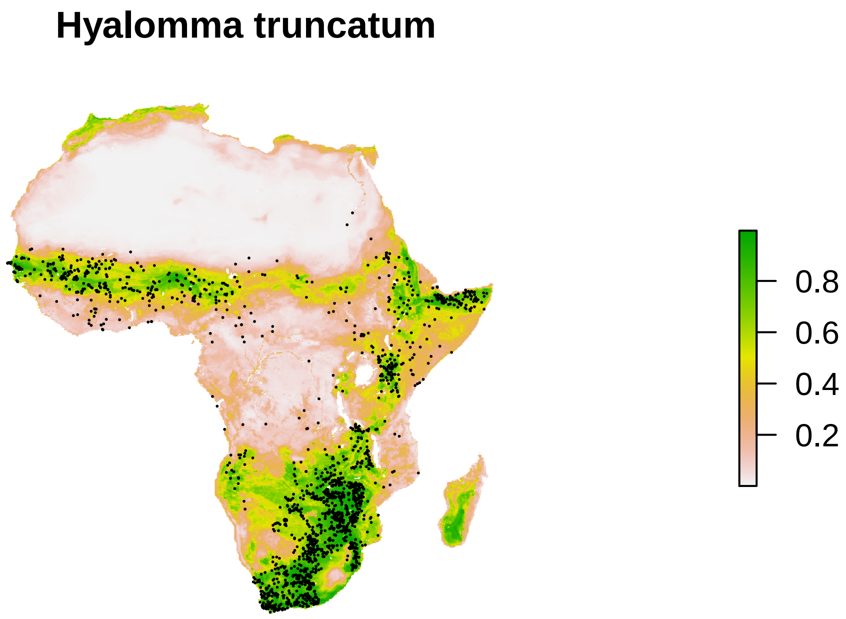


This model seems okay. We're getting predictions in places we don't have any records, like North Africa. That could be good if we think that's suitable climatic space (and if you know *Hyalomma*, you know there's definitely some species there, though posibly not truncatum), but with much of the inhabited area not being predicted, let's revisit that later.

#### Mapping CCHF

Alright. Now let's get back to business by building the CCHF map. We're going to use the same predictors as we used for *H. truncatum*, plus the suitability layer for the ticks. (Using one SDM layer in another map is a sort of finnicky business - sometimes it improves predictions and is epistemologically valid, other times it predetermines a certain outcome in the final model, given issues with colinearity. That's a broader discussion than what we're doing here! If the final map predicts some areas that aren't where the ticks are, I'll call that a win. If they were identical, I would worry.)

##### Running the CCHF model

This time, let's just do the variable selection up front. Same plan as before:

```
# Update the covariate stack

covs <- stack(covs, hytr.layer)
names(covs)[12]='hytr' # for Hyalomma truncatum
xvars <- c(xvars, 'hytr')
```

```r
# Read in the data

load(file = "~/Github/pier39/cchf.rda")
head(cchf)
#>   OCCURRENCE_ID LOCATION_TYPE ADMIN_LEVEL GAUL_AD1 GAUL_AD2
#> 1             1         point        -999     1282    16397
#> 2             2         point        -999     1282    16397
#> 3             3         point        -999     1282    16397
#> 4             4         point        -999     1278    16376
#> 5             5         point        -999     1278    16376
#> 6             6         point        -999     1278    16376
#>   UNIQUE_LOCATION YEAR LATITUDE LONGITUDE    COUNTRY REGION
#> 1             535 1953       38        69 Tajikistan   Asia
#> 2            1178 1953       38        70 Tajikistan   Asia
#> 3             620 1954       42        21     Serbia   Asia
#> 4            1182 1954       37        69 Tajikistan   Asia
#> 5            1165 1954       37        69 Tajikistan   Asia
#> 6            1178 1954       38        70 Tajikistan   Asia
nrow(cchf)
#> [1] 1721

# Spatial thinning checks; this also limits it to African points
# (CCHF is found elsewhere in the world too,
# and our data includes other records!)

cchf <- cchf[,c('LONGITUDE','LATITUDE')]; cchf$Presence = 1
cchf <- SpatialPointsDataFrame(cchf[,1:2],data.frame(Presence=cchf[,3]))
tmp=rasterize(cchf, covs[[1]], field="Presence", fun="min")
pts.sp1=rasterToPoints(tmp, fun=function(x){x>0})
nrow(pts.sp1)
#> [1] 147

# Extract presence values

pres.cov <- raster::extract(covs, pts.sp1[,1:2])
pres.cov <- na.omit(pres.cov)
head(pres.cov)
#>      bio1  bio12 bio13  bio14 bio15 bio2 bio5 bio6  crop ndvi.amp
#> [1,]   31 0.0076 0.011 0.0044  0.20   23   48  9.9 0.046    0.183
#> [2,]   31 0.0076 0.011 0.0044  0.21   23   48  9.9 0.009    0.101
#> [3,]   29 0.0106 0.016 0.0070  0.30   15   41 16.9 0.000    0.031
#> [4,]   31 0.0098 0.015 0.0053  0.33   21   46 14.7 0.036    0.019
#> [5,]   32 0.0085 0.014 0.0036  0.38   21   47 14.5 0.002    0.023
#> [6,]   31 0.0082 0.015 0.0032  0.43   20   47 13.3 0.048    0.025
#>      ndvi.mean  hytr
#> [1,]     0.033 0.021
#> [2,]     0.023 0.034
#> [3,]     0.069 0.177
#> [4,]     0.133 0.381
#> [5,]     0.124 0.126
#> [6,]     0.133 0.187

#Generate pseudoabsences
```

```r
absence <- randomPoints(covs,nrow(pres.cov))
#> Warning in .couldBeLonLat(x, warnings = warnings): CRS is NA. Assuming it
#> is longitude/latitude
abs.cov <- raster::extract(covs, absence)

#Code the response

pres.cov <- data.frame(pres.cov); pres.cov$cchf <- 1
abs.cov <- data.frame(abs.cov); abs.cov$cchf <- 0

# And one to bind them

all.cov <- rbind(pres.cov, abs.cov)
all.cov <- all.cov[complete.cases(all.cov),]; nrow(all.cov)
#> [1] 184
head(all.cov)
#>   bio1  bio12 bio13  bio14 bio15 bio2 bio5 bio6  crop ndvi.amp ndvi.mean
#> 1   31 0.0076 0.011 0.0044  0.20   23   48  9.9 0.046    0.183     0.033
#> 2   31 0.0076 0.011 0.0044  0.21   23   48  9.9 0.009    0.101     0.023
#> 3   29 0.0106 0.016 0.0070  0.30   15   41 16.9 0.000    0.031     0.069
#> 4   31 0.0098 0.015 0.0053  0.33   21   46 14.7 0.036    0.019     0.133
#> 5   32 0.0085 0.014 0.0036  0.38   21   47 14.5 0.002    0.023     0.124
#> 6   31 0.0082 0.015 0.0032  0.43   20   47 13.3 0.048    0.025     0.133
#>    hytr cchf
#> 1 0.021    1
#> 2 0.034    1
#> 3 0.177    1
#> 4 0.381    1
#> 5 0.126    1
#> 6 0.187    1

# This part automates the variable selection and returns the model

cchf.model <- bart.step(x.data = all.cov[,xvars],
                        y.data = all.cov[,'cchf'],
                        full = TRUE,
                        quiet = TRUE)
#>
#>  10 tree models: 100 iterations
#>
#>  20 tree models: 100 iterations
#>
#>  50 tree models: 100 iterations
#>
#>  100 tree models: 100 iterations
#>
#>  150 tree models: 100 iterations
#>
#>  200 tree models: 100 iterations
```

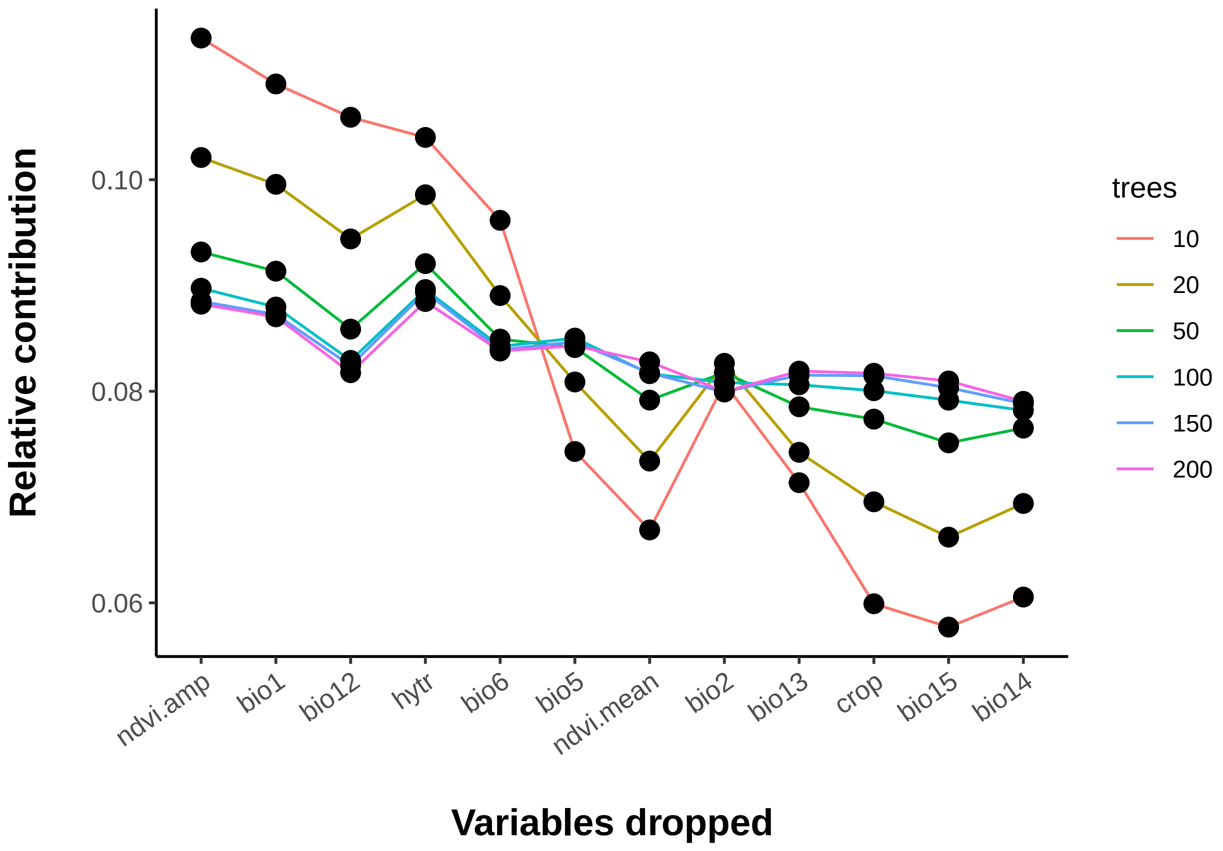


```
#> [1] Number of variables included: 12
#> [1] Dropped:
#> [1]
#> [1] ---------------------------------------
#> [1] Number of variables included: 11
#> [1] Dropped:
#> [1] bio15
#> [1] ---------------------------------------
#> [1] Number of variables included: 10
#> [1] Dropped:
#> [1] bio15 crop
#> [1] ---------------------------------------
#> [1] Number of variables included: 9
#> [1] Dropped:
#> [1] bio15 crop  bio14
#> [1] ---------------------------------------
#> [1] Number of variables included: 8
#> [1] Dropped:
#> [1] bio15     crop      bio14     ndvi.mean
#> [1] ---------------------------------------
#> [1] Number of variables included: 7
#> [1] Dropped:
#> [1] bio15     crop      bio14     ndvi.mean bio13
#> [1] ---------------------------------------
#> [1] Number of variables included: 6
#> [1] Dropped:
```

```
#> [1] bio15     crop      bio14     ndvi.mean bio13     bio2
#> [1] ---------------------------------------
#> [1] Number of variables included: 5
#> [1] Dropped:
#> [1] bio15     crop      bio14     ndvi.mean bio13     bio2      bio5
#> [1] ---------------------------------------
#> [1] Number of variables included: 4
#> [1] Dropped:
#> [1] bio15     crop      bio14     ndvi.mean bio13     bio2      bio5
#> [8] hytr
#> [1] ---------------------------------------
#> [1] Number of variables included: 3
#> [1] Dropped:
#> [1] bio15     crop      bio14     ndvi.mean bio13     bio2      bio5
#> [8] hytr      bio1
#> [1] ---------------------------------------
```

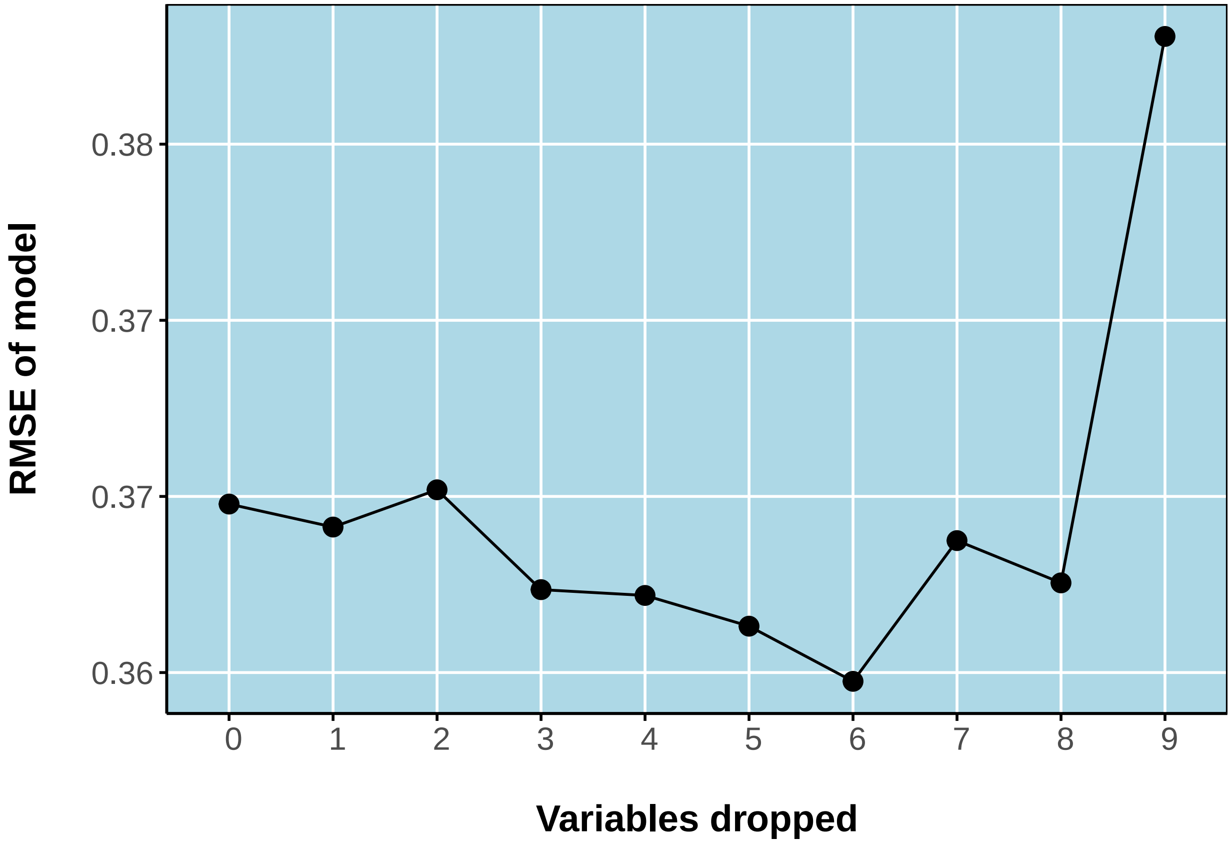


```
#> [1] ---------------------------------------
#> [1] Final recommended variable list
#> [1] bio1     bio12    bio5     bio6     ndvi.amp hytr
#>
#> Running BART with binary y
#>
#> number of trees: 200
#> number of chains: 1, number of threads 1
#> Prior:
#>  k: 2.000000
```

```
#>  power and base for tree prior: 2.000000 0.950000
#>  use quantiles for rule cut points: false
#> data:
#>  number of training observations: 184
#>  number of test observations: 0
#>  number of explanatory variables: 6
#>
#> Cutoff rules c in x<=c vs x>c
#> Number of cutoffs: (var: number of possible c):
#> (1: 100) (2: 100) (3: 100) (4: 100) (5: 100)
#> (6: 100)
#>
#> offsets:
#>  reg : 0.00 0.00 0.00 0.00 0.00
#> Running mcmc loop:
#> iteration: 100 (of 1000)
#> iteration: 200 (of 1000)
#> iteration: 300 (of 1000)
#> iteration: 400 (of 1000)
#> iteration: 500 (of 1000)
#> iteration: 600 (of 1000)
#> iteration: 700 (of 1000)
#> iteration: 800 (of 1000)
#> iteration: 900 (of 1000)
#> iteration: 1000 (of 1000)
#> total seconds in loop: 1.149624
#>
#> Tree sizes, last iteration:
#> [1] 4 2 3 2 2 2 2 2 2 2 6 2 3 2 1 2 3 2
#> 2 2 1 2 5 3 2 4 2 2 3 3 3 2 2 2 4 3 2 3
#> 1 2 4 4 1 4 3 2 2 3 4 3 3 2 3 2 3 2 2 1
#> 2 3 2 2 2 2 3 3 6 2 3 2 2 2 2 2 4 2 4 2
#> 2 2 3 3 4 2 4 2 2 2 3 2 2 2 2 2 2 4 2 2
#> 3 3 3 4 1 2 2 2 2 1 2 3 2 3 3 2 2 2 2 1
#> 3 4 3 2 2 3 2 2 2 3 2 4 2 3 2 2 2 2 2 3
#> 2 2 2 2 2 2 2 2 2 2 2 2 2 2 3 1 1 3 3 3
#> 3 2 2 2 2 2 2 2 2 2 2 1 2 3 2 2 2 2 2 2
#> 2 2 2 2 4 2 2 3 3 2 4 2 2 2 2 5 4 3 2 2
#> 3 2
#>
#> Variable Usage, last iteration (var:count):
#> (1: 50) (2: 50) (3: 54) (4: 39) (5: 46)
#> (6: 44)
#> DONE BART
```

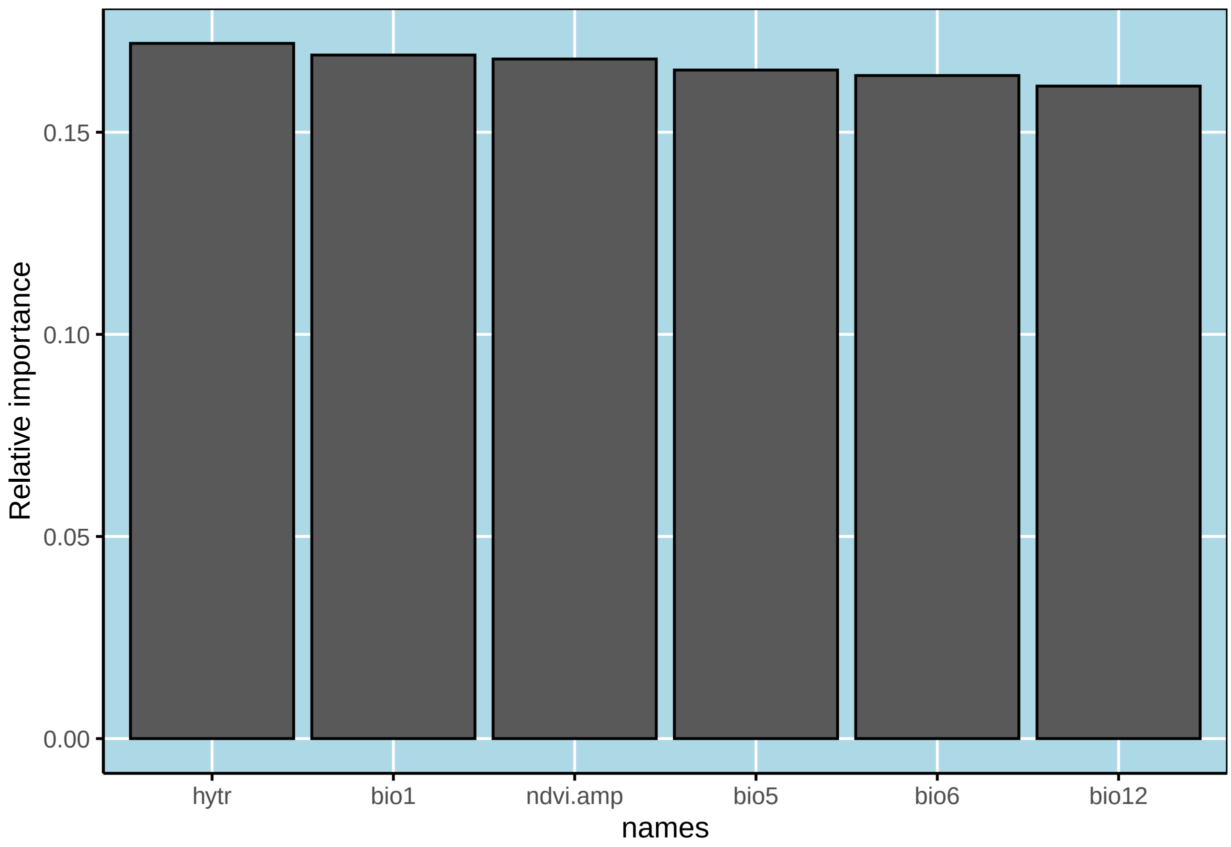


```
#> Call:  bart x.data[, vs] y.data TRUE
#>
#> Predictor list:
#>  bio1 bio12 bio5 bio6 ndvi.amp hytr
#>
#> Area under the receiver-operator curve
#> AUC = 0.91
#>
#> Recommended threshold (maximizes true skill statistic)
#> Cutoff =  0.47
#> TSS =  0.66
#> Resulting type I error rate:  0.15
#> Resulting type II error rate:  0.18
```

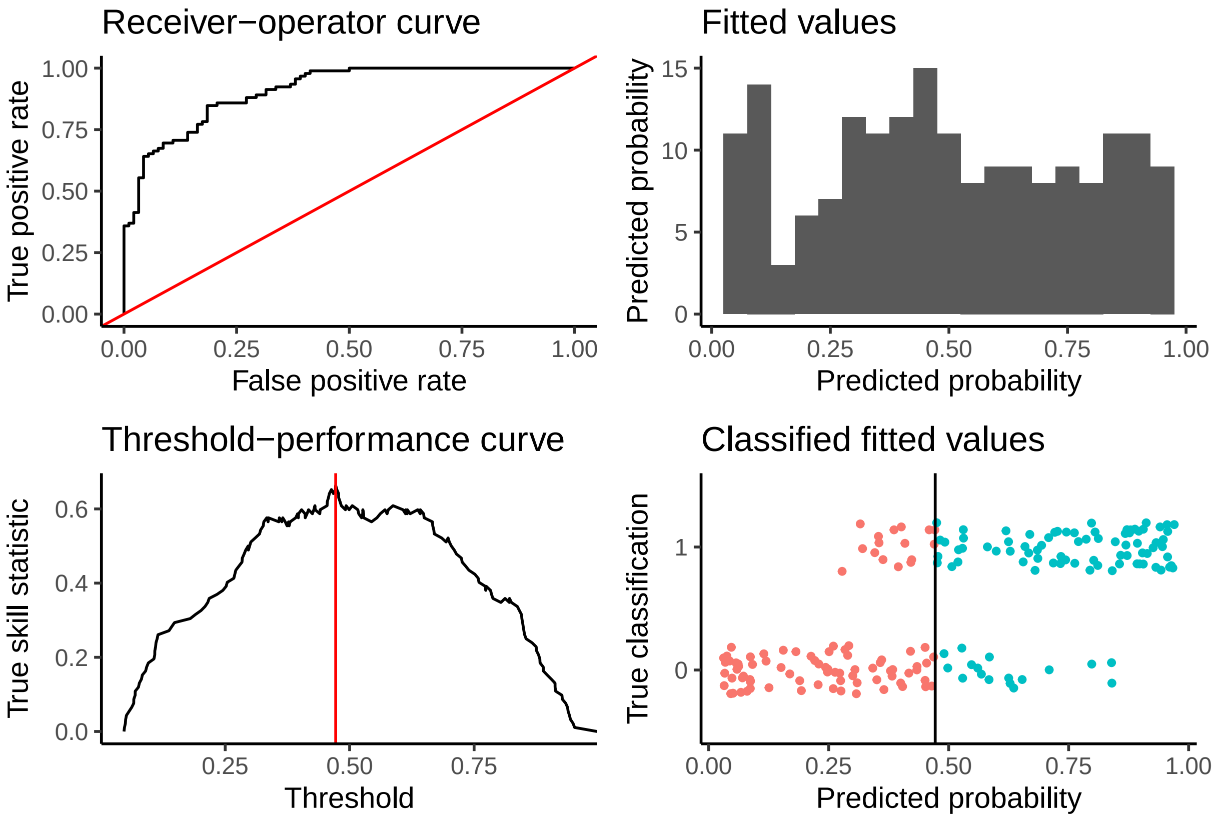


```
# Do the spatial prediction
# This time, let's add the quantiles

cchf.map <- predict(object = cchf.model,
                    inputstack = covs,
                    quantiles=c(0.025, 0.975),
                    splitby=20,
                    quiet = TRUE)
#> Estimated time to total prediction (mins):
#> 8.4

# How's it look?
plot(cchf.map[[1]], box=FALSE, axes=FALSE, main='CCHF')
points(pts.sp1, col='black',
        pch=16, cex=0.2)
```

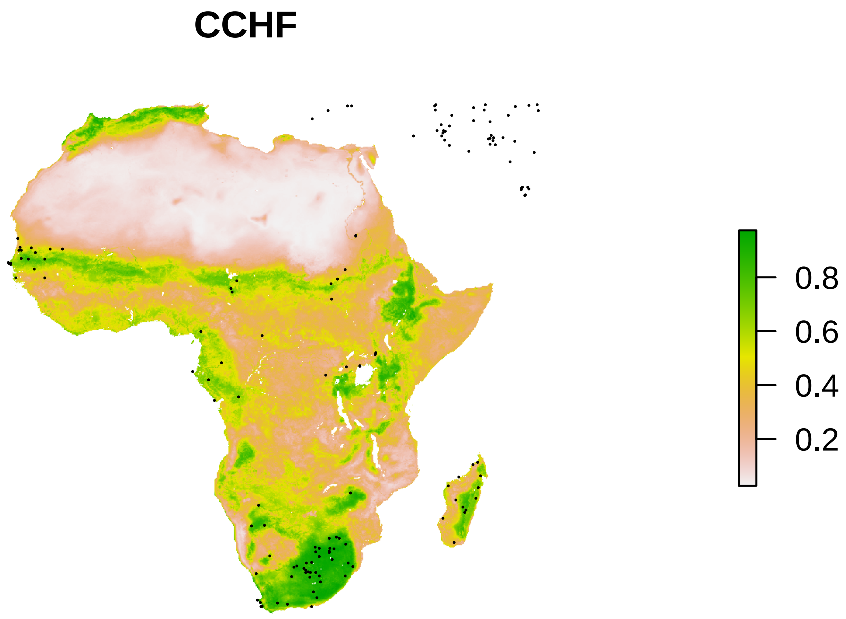


OK. Nice model! Let's see what we can do to unpack it.

First, let's compare it against the tick map.

```
par(mfrow=c(1,2))
plot(hytr.layer, box=FALSE, axes=FALSE, main='H. truncatum')
plot(cchf.map[[1]], box=FALSE, axes=FALSE, main='CCHF')
```

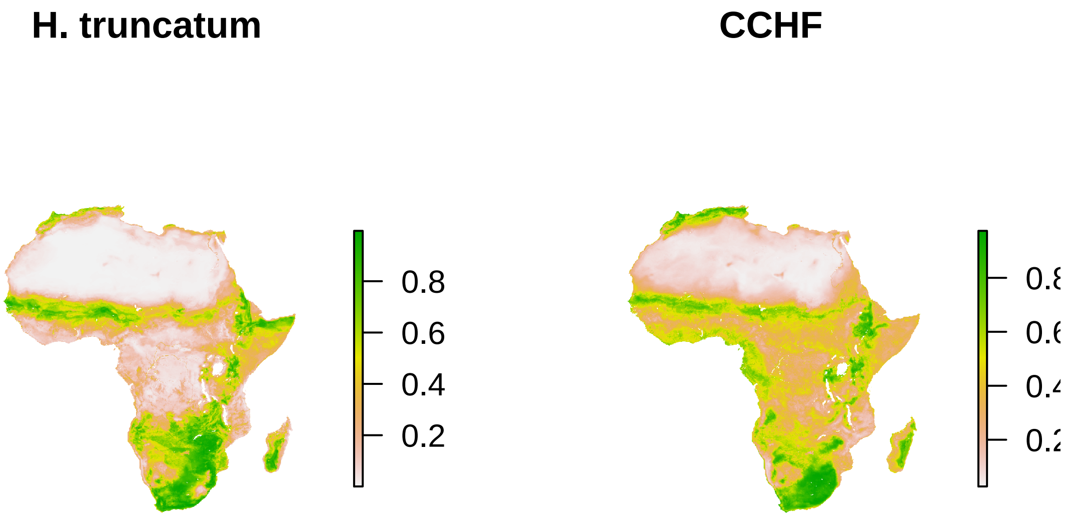


Our model seems to be different from the previous one (published by Messina *et al.*, using this dataset which they generously provide online) in three major ways.

1. First, using the tick vector has increased the amount of predicted suitable area in South Africa, Namibia, Botswana, and Zimbabwe. That makes sense, overall–if there are vectors present, CCHF seems plausible.
2. The model predicts the area in coastal Cameroon, Gabon, and Equatorial Guinea that we know has some CCHF records but has previously been underpredicted. Weirdly, we don't have good evidence *Hyalomma truncatum* is there. So, the model is doing well, but the ecology is still unclear.
3. Finally, the northern coast of Africa is predicted to be highly suitable. There's plenty of *Hyalomma* species up there, though not *H. truncatum* as far as our data suggests. It's possible we should think more about the possibility of CCHF in Morocco and Algeria.

Let's take a look at the uncertainty in the model:

```
par(mfrow=c(1,2))
plot(cchf.map[[2]], box=FALSE, axes=FALSE, main='2.5% bound')
plot(cchf.map[[3]], box=FALSE, axes=FALSE, main='97.5% bound')
```

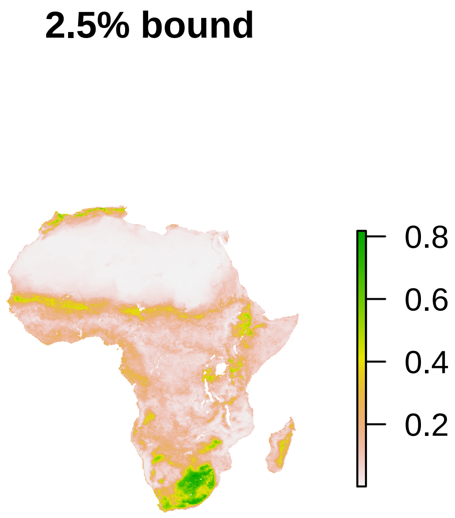


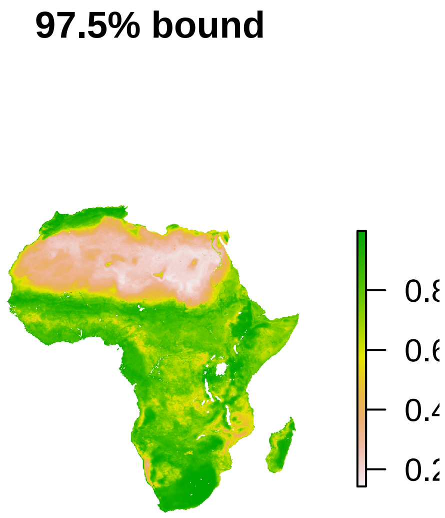


This may be more interesting to think about in the context of binary maps of presence/absence. We can produce those using `summary`:

```
summary(cchf.model)
#> Call:  bart x.data[, vs] y.data TRUE
#>
#> Predictor list:
#>  bio1 bio12 bio5 bio6 ndvi.amp hytr
#>
#> Area under the receiver-operator curve
#> AUC = 0.91
#>
#> Recommended threshold (maximizes true skill statistic)
#> Cutoff =  0.47
#> TSS =  0.66
#> Resulting type I error rate:  0.15
#> Resulting type II error rate:  0.18
```

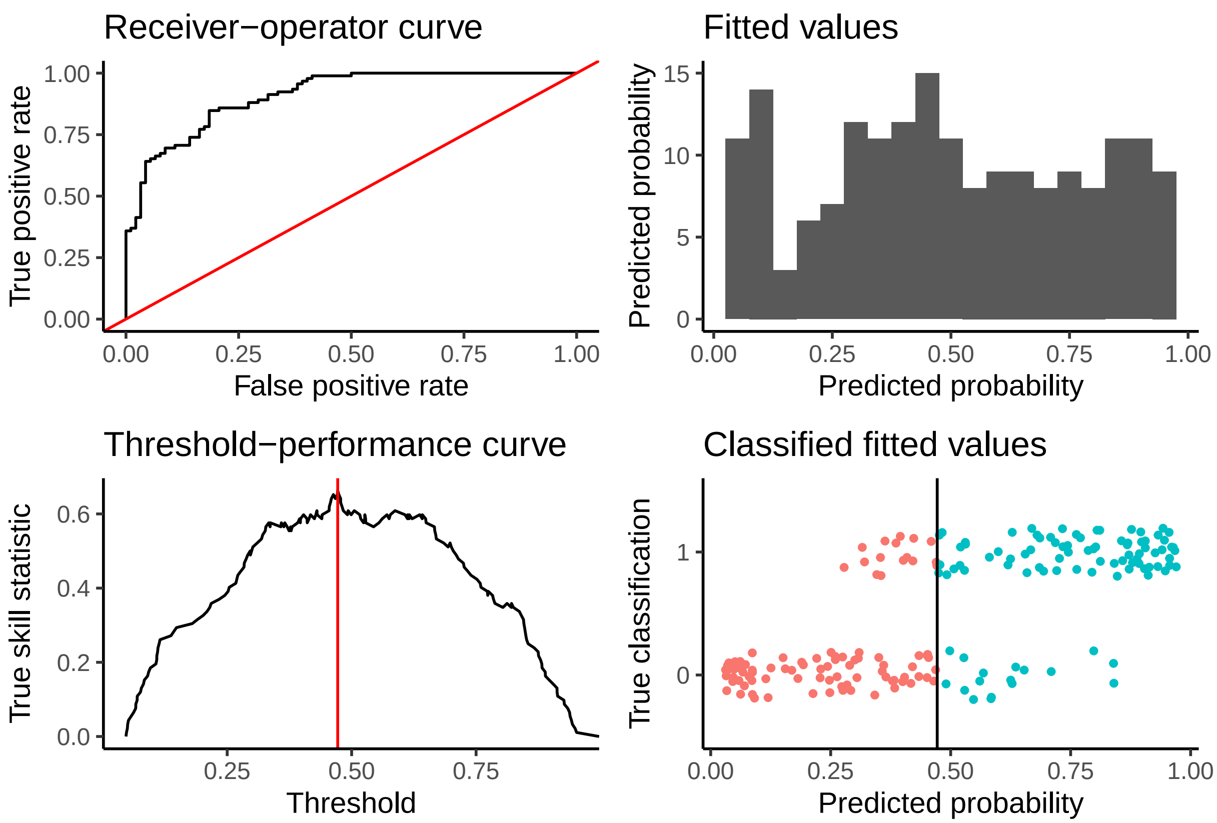


```
par(mfrow=c(1,1))
plot(cchf.map[[1]]>0.47, box=FALSE, axes=FALSE, main='CCHF risk')
```

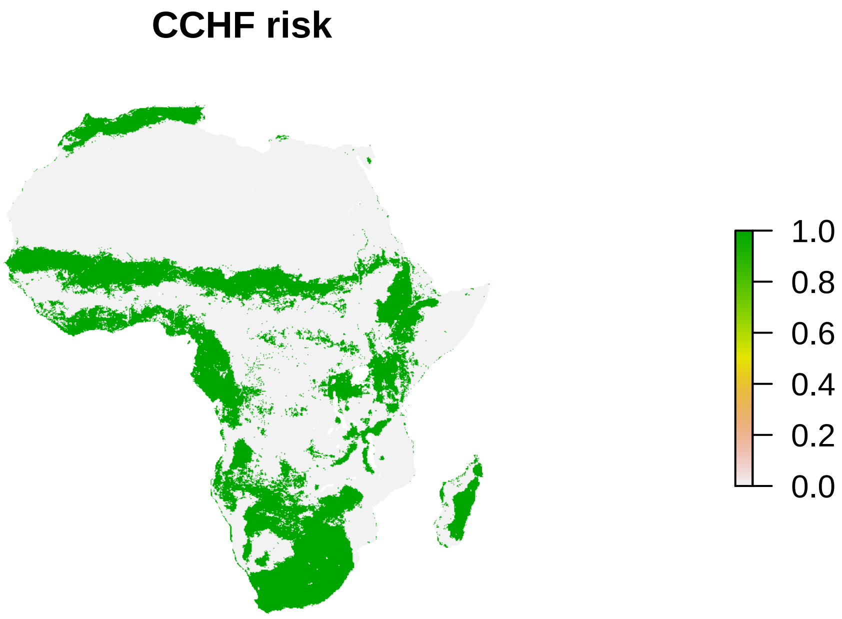


```
par(mfrow=c(1,2))
plot(cchf.map[[2]]>0.47, box=FALSE, axes=FALSE, main='2.5% bound')
plot(cchf.map[[3]]>0.47, box=FALSE, axes=FALSE, main='97.5% bound')
```

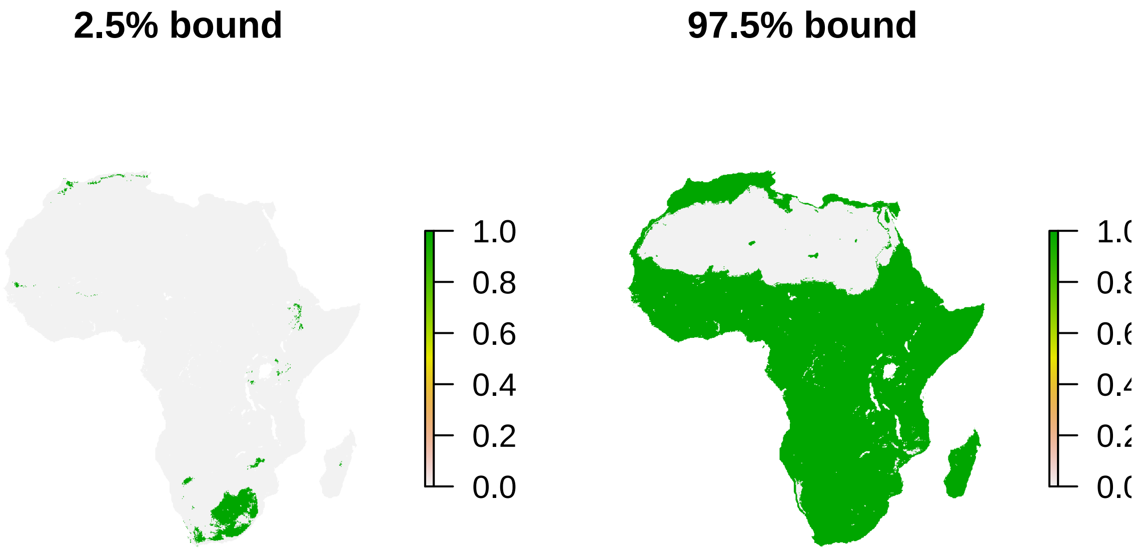


The outer bounds here are *incredibly* wide - which highlights exactly how little data we have about the distribution of CCHF, and how much uncertainty presentation matters.

Where is the uncertainty highest?

```
par(mfrow=c(1,1))
plot(cchf.map[[3]]-cchf.map[[2]],
     box=FALSE, axes=FALSE, main='Posterior width, scaled')
```

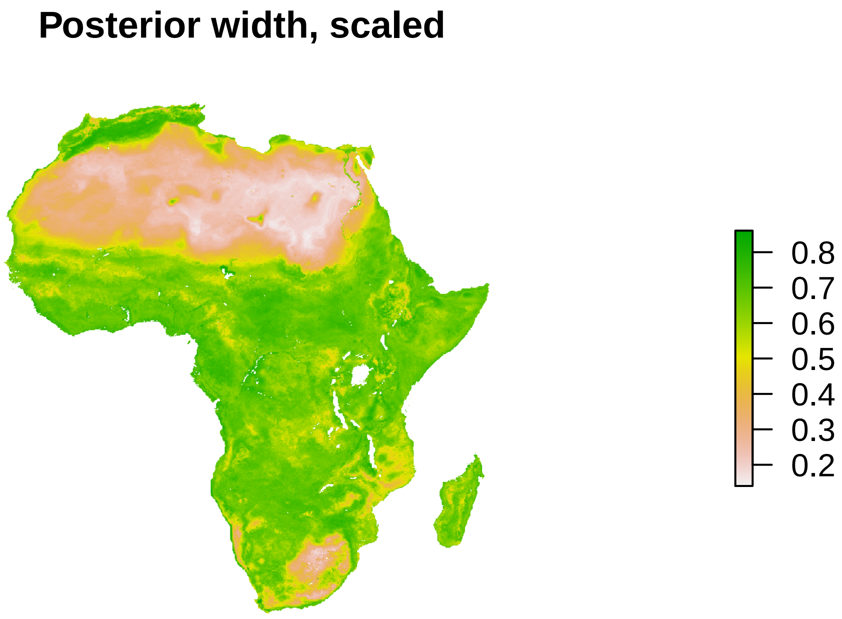


Posterior width isn't always the most informative measure of uncertainty. But it does tell us we're most confident in the Sahara, where we're quite sure there's no real CCHF risk, and in the parts of South Africa where spillover is most common. We can further investigate by mapping the places with the most uncertainty:

```
quant <- quantile(values(cchf.map[[3]]-cchf.map[[2]]), 0.75, na.rm=TRUE)

par(mfrow=c(1,1))
plot((cchf.map[[3]]-cchf.map[[2]])>quant,
     box=FALSE, axes=FALSE,
     main = "Highest uncertainty zones")
```

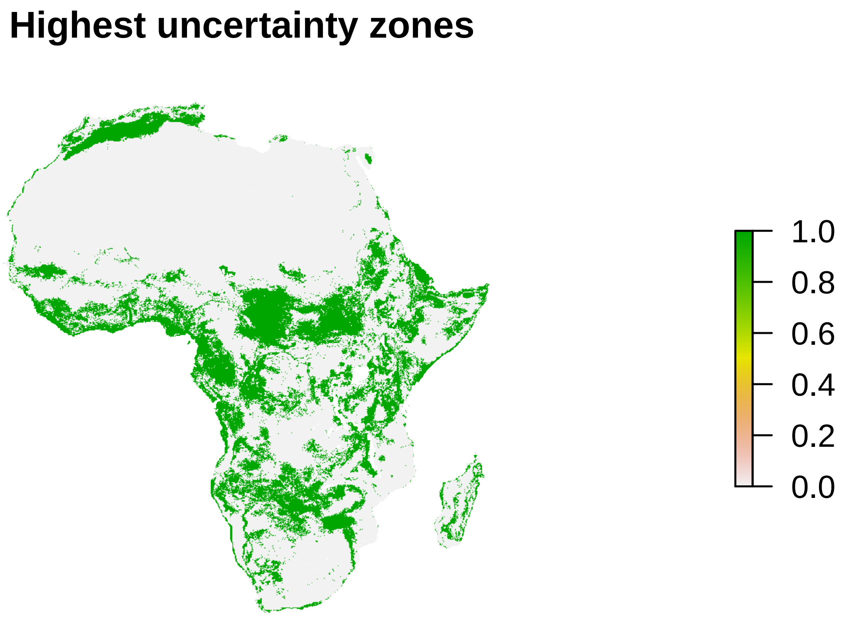


The most interesting takehome is that uncertainty is particularly high in the Sahel and on the Western coast of Africa; along the equator, CCHF spillover has previously occurred but the tick vector is absent. This may be worth further investigation. Similarly, there are no spillovers along the southern coast of West Africa, but uncertainty is very high, and it may be worth investigating this more (especially given tick records in that region).

#### Analytics

Finally, let's unpack some of what's under the hood in the model. First let's look at the variable contributions:

```
varimp(cchf.model, plots=TRUE)
```

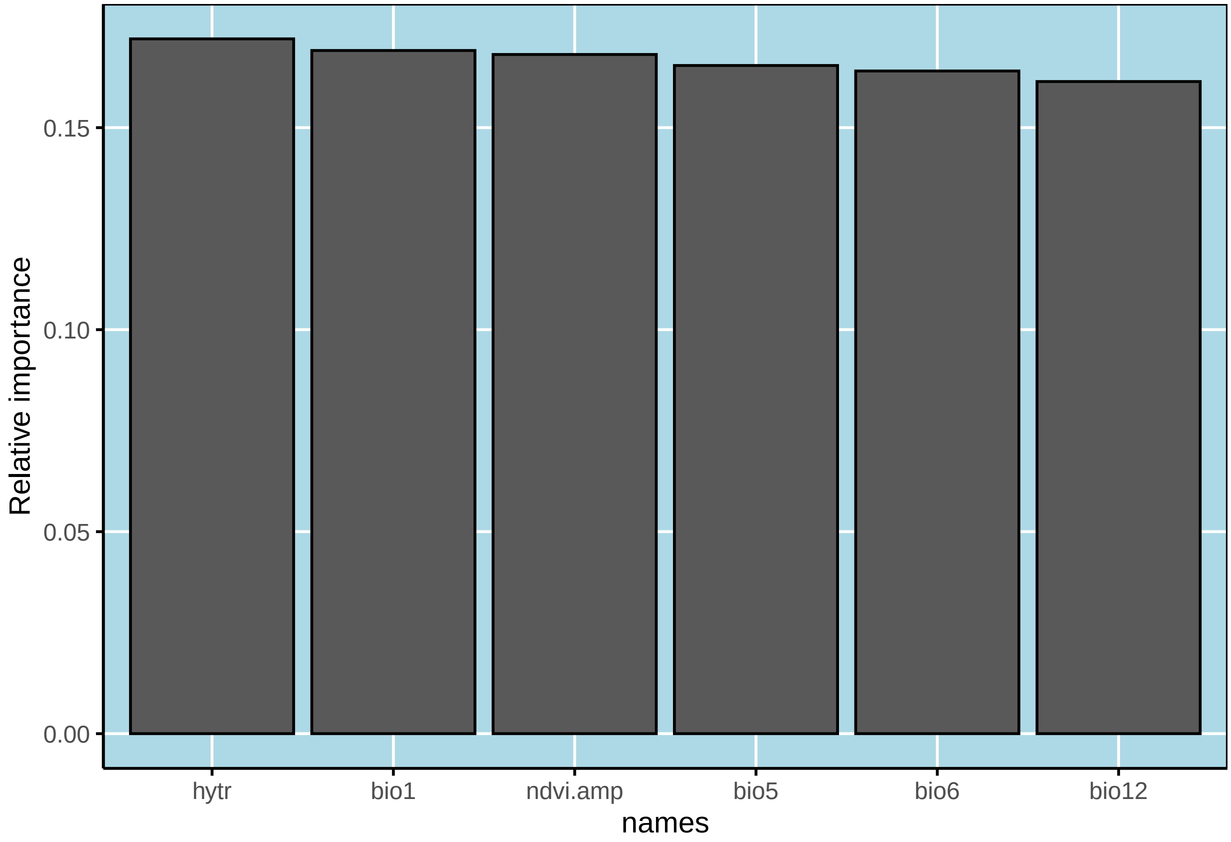


```
#>      names varimps
#> 1     bio1    0.17
#> 2    bio12    0.16
#> 3     bio5    0.17
#> 4     bio6    0.16
#> 5 ndvi.amp    0.17
#> 6     hytr    0.17
```

The tick vector comes out on top, unsurprisingly! Let's look at the response functions a little bit. First, let's look at the partial dependence plots for a couple individual variables.

```
# Let's check the tick response curve
p = partial(cchf.model,
        'hytr',
        trace=FALSE,
        ci=TRUE,
        equal=TRUE,
        smooth=3)
```

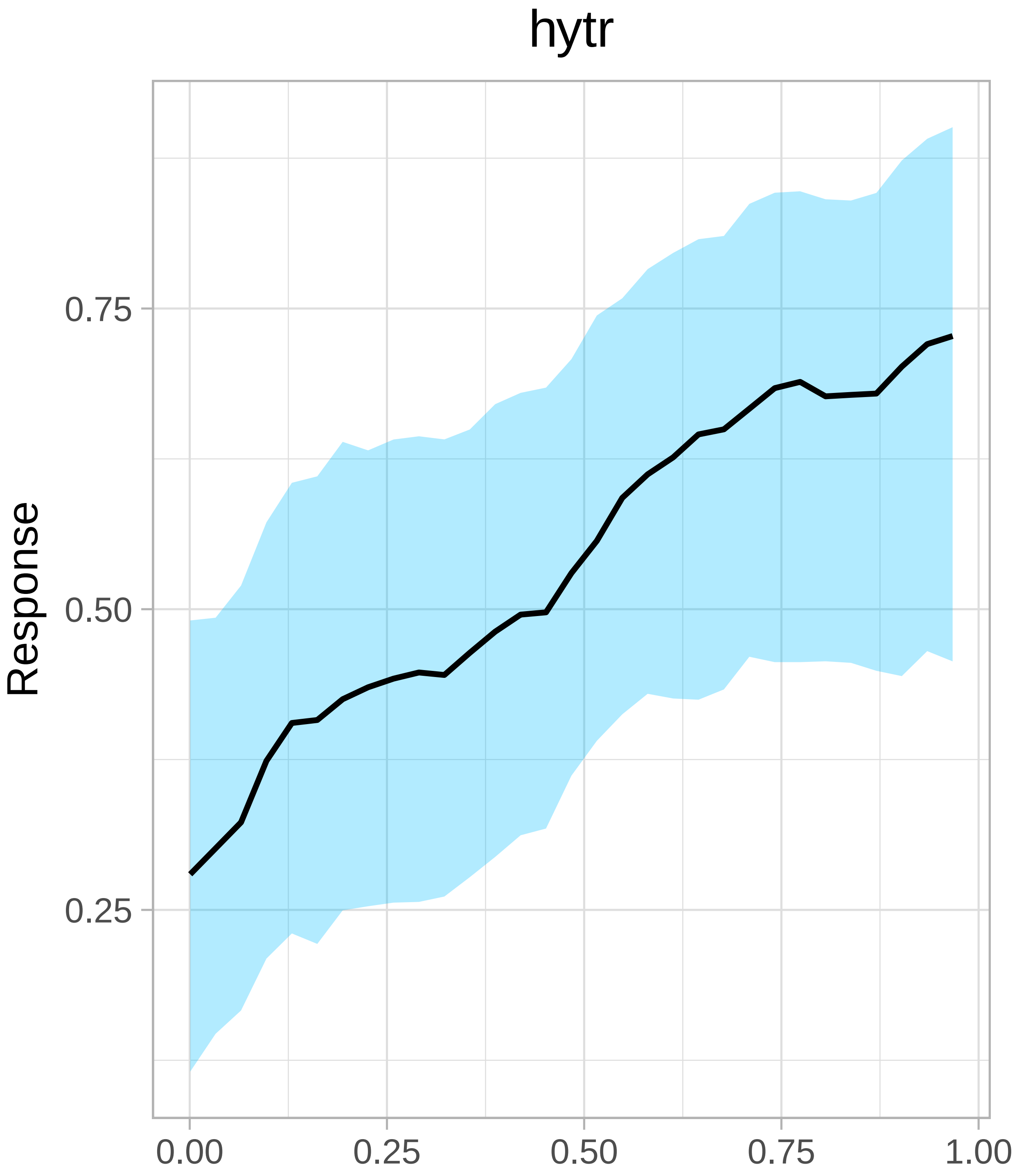


```
# Let's do a few at once
partial(cchf.model,
        c('bio1','bio5','bio6','bio12'),
        trace=FALSE,
        ci=TRUE,
```

```
        panel=TRUE,
        smooth=3)
```

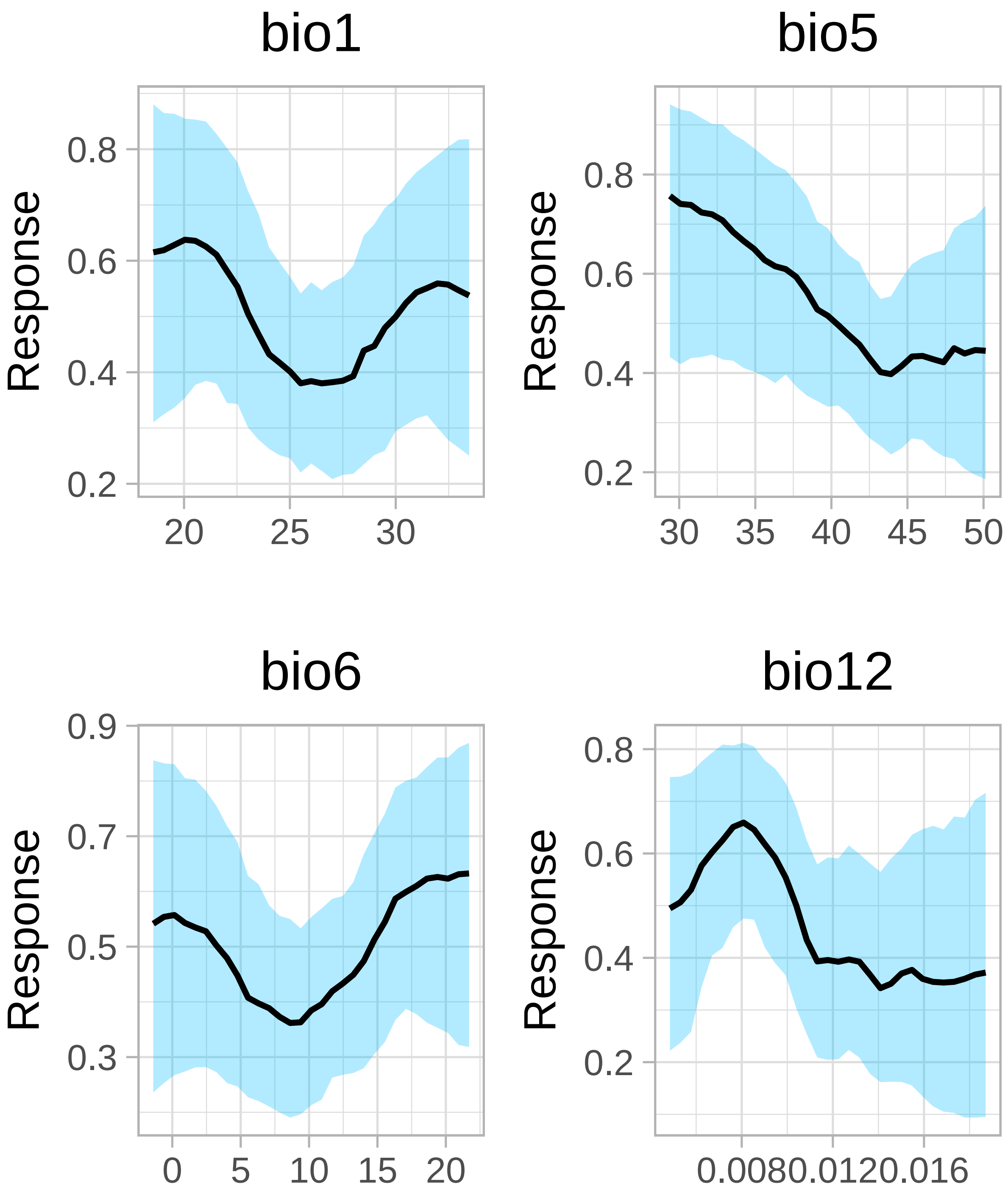


Some of these patterns are pretty clear - suitability declines with the maximum temperature of the warmest month, and increases with the probability of the tick.

A great feature of BART is that we can pretty easily do two-dimensional partial dependence plots, and we can pretty easily visualize the optimum (I haven't added a wrapper for this yet):

```
p <- pd2bart(cchf.model, xind=c('bio12', 'ndvi.amp'), pl=TRUE)
```

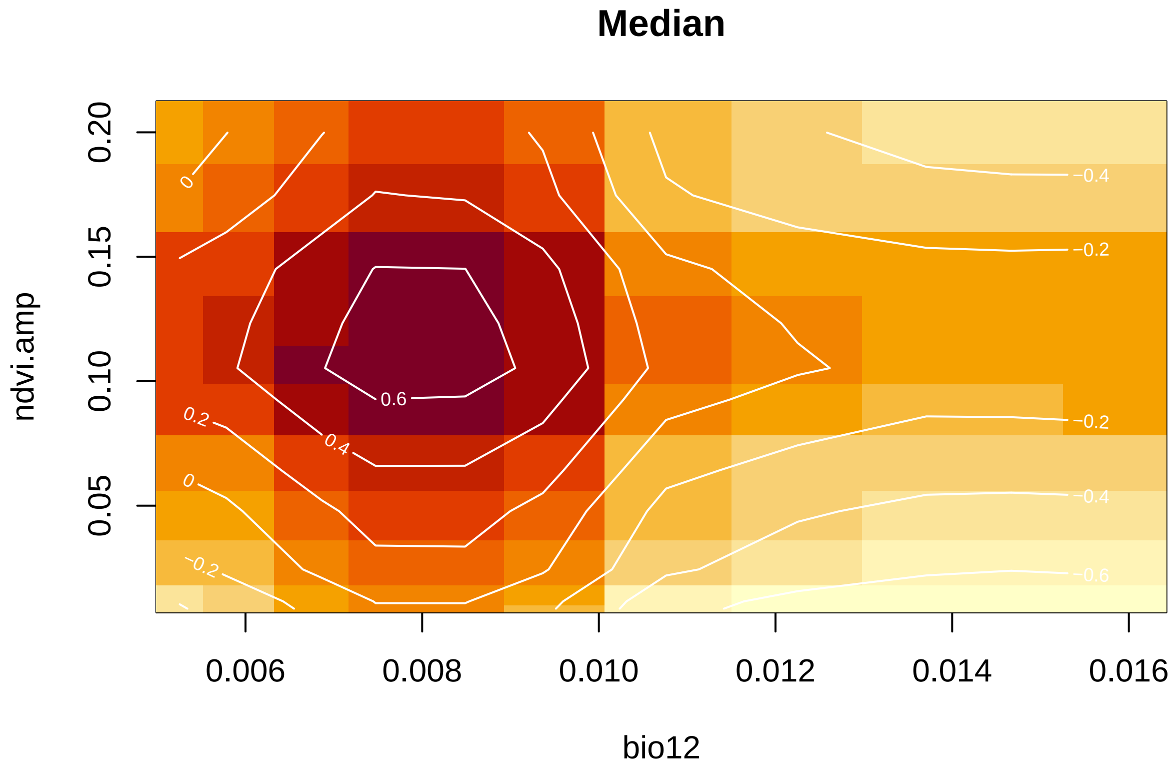


This is probably my favorite feature of `dbarts` - these plots are a really nice way to visualize the Hutchinsonian niche sort of like how NicheA does, but within the familiar framework of classification trees. Plus it's not just a cross-product of the individual partials - the BART framework allows for interactions and sometimes you'll see them show up in these plots.

One last cool trick: spatial partial dependence plots

```
spartial(cchf.model, covs, x.vars = 'bio1', equal=TRUE)
```

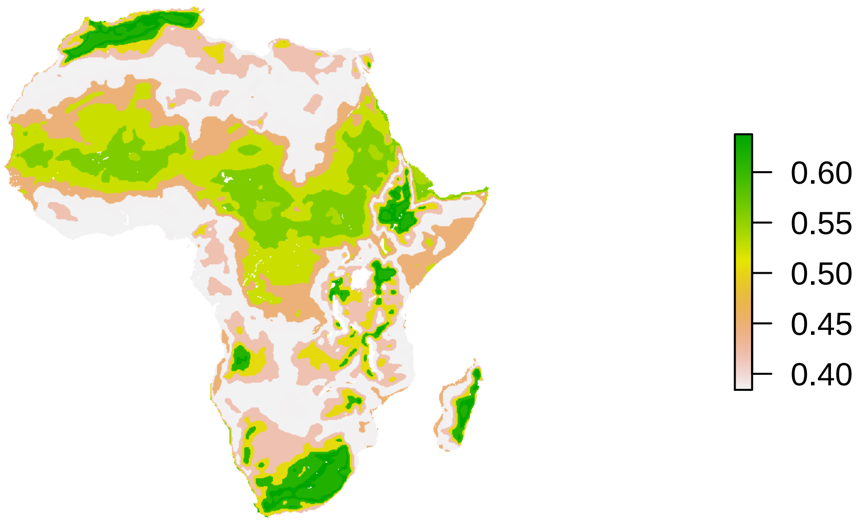


```
#> class       : RasterStack
#> dimensions : 867, 999, 866133, 1  (nrow, ncol, ncell, nlayers)
#> resolution : 0.083, 0.083  (x, y)
#> extent     : -25, 58, -35, 37  (xmin, xmax, ymin, ymax)
#> crs        : NA
#> names      : bio1
#> min values : 0.38
#> max values : 0.64
```

This map projects the partial dependence plot onto the raster data for the bio1 layer, and can be interpreted as answering the question "Where are mean temperatures most conducive for viral transmission?"

#### Finally…

A big thanks to Jane Messina *et al.* for publicly sharing their CCHF data, and to Graeme Cumming for tick data.
